# iFLiP: accessible and low-cost lifetime fiber photometry for robust longitudinal and quantitative measurements of neural dynamics

**DOI:** 10.64898/2026.10.21.760747

**Authors:** Takaki Yahiro, Yang Chen, Landon Bayless-Edwards, Sophia An, Tianyi Mao, Haining Zhong

## Abstract

Fluorescence lifetime measurements overcome the limitation of widely used intensity-based measurements in quantifying tonic activities and longitudinal changes in neural dynamics. However, while lifetime sensors are rapidly emerging, their applications have been limited by the complexity and expense of required hardware and software. Here, we report a simple, low-cost lifetime fiber photometry system, named iFLiP, that uses entirely off-the-shelf products and can be assembled without specialized expertise. We also develop software that enables turn-key operation. The integrated system can accurately monitor lifetime changes at up to 5000 Hz for days. We demonstrate the utility of iFLiP in monitoring the activity of protein kinase A (PKA) in vitro and in vivo, and uncover striatal subregion– and cell-type-specific PKA dynamics elicited by locomotion in mice. Additionally, multi-day continuous measurements in the basal forebrain reveal previously unappreciated PKA activity associated with sleep/wakefulness. Overall, iFLiP may make lifetime measurements broadly accessible in neuroscience research.

## INTRODUCTION

Brain function relies on its underlying neural activity. Genetically encoded sensors, such as those for calcium, voltage, neurotransmitters, and intracellular signaling events, have begun to enable measurements of diverse neural activities in living animals.^1–3^ To date, most studies have relied on intensity-based sensors, which excel in monitoring short-term, relative changes. However, long-term changes and tonic analyte levels are also critical determinants of cellular, circuit, and behavioral function. Yet, quantifying these dynamics remains challenging because the intensity measurements are confounded by common variables, such as light scattering, signal or sample drifting, photobleaching, and sensor expression levels.

There is growing recognition that these limitations may be overcome by measuring fluorescence lifetime changes of appropriate sensors. The fluorescence lifetime of a fluorophore is independent of the fluorescence intensity and therefore insensitive to the aforementioned confounding factors.^4–6^ Lifetime measurements have been used to monitor longitudinal analyte changes, compare absolute signaling activities across samples or recording sessions, and improve the signal-to-noise ratio in vitro and in vivo (e.g., ref^7–14^). At present, lifetime sensors are becoming increasingly available.^9,12,13,15–18^ However, despite previous efforts, fluorescence lifetime measurements remain inaccessible to most laboratories due to the high cost, complicated software and hardware, and specialized expertise required.

Towards making the technology more accessible, lifetime measurement has been integrated with fiber photometry.^15,19^ The first fluorescence lifetime photometry (FLiP) system used the time-domain measurement principle, which is advantageous under low photon flux conditions.^19^ Mechanistic insights―different lifetimes of two sensor states (sensor properties) and their relative fractions (biologically-relevant readout)―can be obtained from a single measurement. However, this system required both optical engineering expertise and expensive components. Additionally, this system had a limited sampling rate of ∼1 Hz and was prone to errors even at moderate signal intensities (>1×10^6^ photons/s). Another FLiP system, FLIPR, utilized the frequency-domain measurement principle.^15^ While it improved upon many issues of the first system, it also required advanced expertise in both optics and electronics, making it challenging to adopt in typical biological laboratories. The system also remained relatively expensive (∼$38k). Importantly, with a single stimulation frequency using the frequency-domain measurement, the system may encounter difficulties in dissecting the contributions of different lifetimes versus the relative abundance of multiple lifetime populations.^20–22^ As a result, while intensity-based fiber photometry has proven to be accessible to many laboratories, the current FLiP systems remain challenging to adopt and apply.

Here, we develop iFLiP, a simple, low-cost, yet flexible and robust system for lifetime fiber photometry. This system uses entirely off-the-shelf components and can be assembled using standard connectors and cables. It is alignment-free, requiring no specialized expertise in optics or electronics. Once assembled, our custom-developed software enables turn-key data acquisition and streamlined analysis. By employing strategies to eliminate the “pile-up” effect, which causes systematic errors at high intensities, and by optimizing single-photon data processing, iFLiP can monitor lifetime at sampling rates up to 5000 Hz and intensities of up to 2×10^7^ photons/s. We demonstrated the utility of iFLiP across multiple preparations, including organic dyes, cultured cells, acute brain slices, and in vivo recordings in behaving animals. Using a well-characterized lifetime sensor for protein kinase A (PKA), which plays a central role in mediating neuromodulatory regulation, we found that animal locomotion elicited PKA dynamics with subregion– and cell-type-specific characteristics in the striatum. Additionally, continuous recording over days in the basal forebrain revealed novel dynamics of PKA activity that were associated with locomotion and circadian rhythms. Taken together, these results establish iFLiP as an accessible platform for quantitative and longitudinal fluorescence lifetime measurements of brain activity.

## RESULTS

### Design of iFLiP

To develop iFLiP, we chose to use the time-domain measurement for two reasons: i) its sensitivity under low-light conditions, which are common in biological samples, and ii) its ability to quantify, from a single measurement, the relative populations of different lifetime components for mechanistic dissection.

To eliminate the need for specialized expertise and minimize complexity, we optimized each key component (Figure 1A). First, we chose gain-switched, picosecond-pulsed diode lasers, which are inexpensive and available in fiber-coupled formats.^23^ Second, we used modern fiber optics that enable the use of filter cubes without mirror-based optical alignment. Third, we designed a simplified light path so that the emission light returning from the sample is routed through the same filter cube into a fiber that is fed directly to a detector without collection lenses. Effective light collection can be achieved by positioning the fiber close to a detector with a relatively large active area (ϕ > 1 mm). Fourth, we selected a low-cost, sensitive (quantum yield ∼40%), temperature-compensated silicon photomultiplier (SiPM) detector, which features fast response kinetics for single-photon measurements, high resistance to strong light exposure, and a sufficiently large active area.

**Figure 1.**
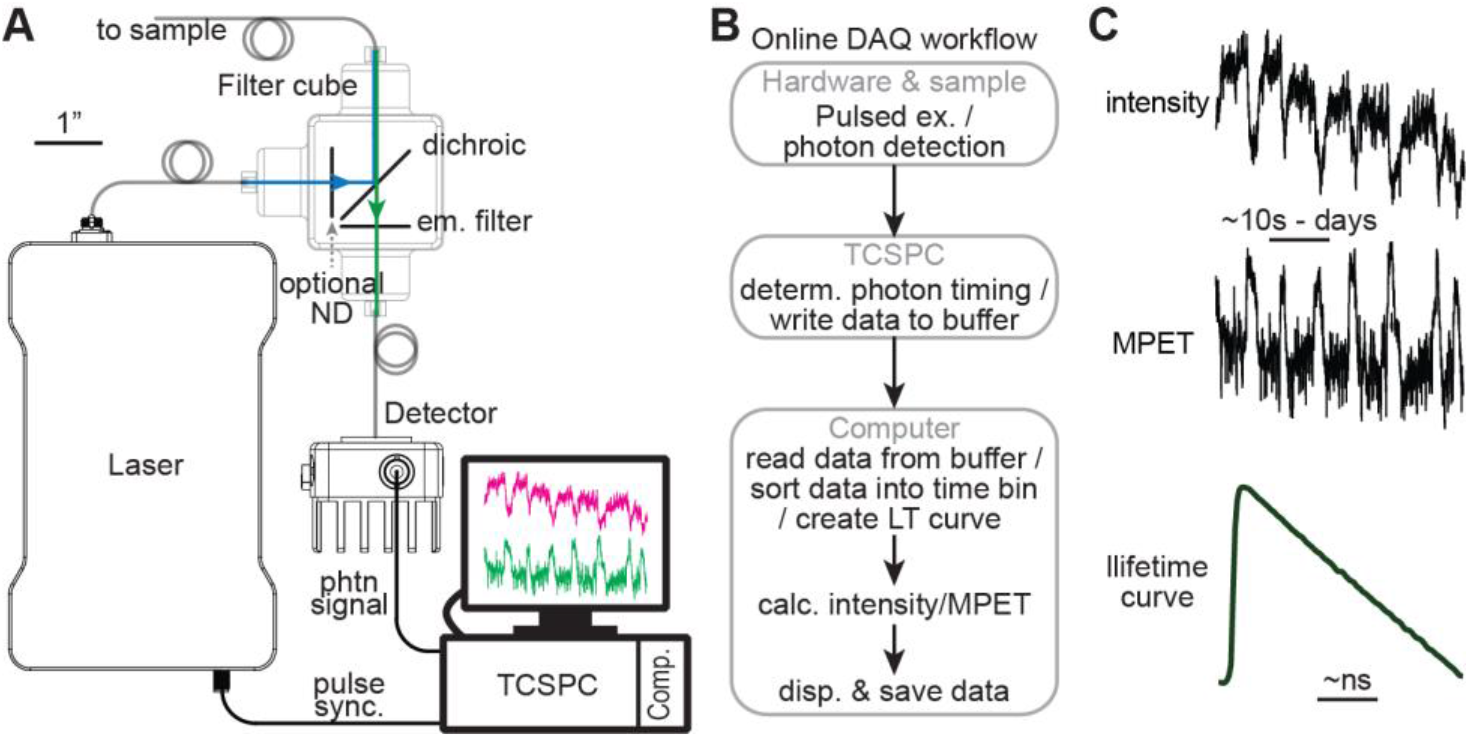
Conceptual design of iFLiP. (A) Schematic illustrating the key components of the iFLiP hardware. (B) Conceptual flowchart of data acquisition and processing. (C) Schematic illustrating the data iFLiP records and displays live: the intensity, MPET, and lifetime curve. Em.: emission, sync.: synchronization, phtn: photon, disp.: display, calc.: calculation. See also Figures S1 – S3.

The timing signal from the laser (i.e., the sync signal) and the photon signal from the detector are fed into a computer board for time-correlated single-photon counting (TCSPC),^4,5,22^ from which the lifetime is determined (Figure 1A). Single-photon signals are identified using a constant fraction discriminator (CFD), which detects the timing of pulses with different amplitudes (due to noise in amplification gains) with high precision.^24^ The CFD triggers a time-to-digital converter (TDC) to digitize photon timing with a shorter deadtime (25 ns) and reduced cost compared to previous systems.

The previous time-domain FLiP system was limited to detecting low signal intensities due to an effect called “pile-up”.^25,26^ This is caused by the deadtime in the TCSPC process: when a photon is detected, a second photon arriving within the deadtime cannot be measured, thereby distorting the probability of photon detection over time at high photon fluxes (Figure S1A and S1B). However, when the inter-pulse interval is an integer fraction of the dead time (i.e., the multiplication of laser frequency and dead time being an integer), the photon detection probability is uniformly affected throughout the inter-pulse interval, thereby eliminating distortion of lifetime measurements (Figure S1A and S1B). Thus, we set the laser frequency at 80 MHz, which corresponds to an inter-pulse interval of 12.5 ns, half of the deadtime of the TCSPC board we used (25 ns). This allows for a greatly increased intensity limit, which is imposed by the loss of photons due to deadtime (Figure S1C).

In parallel, we developed software with a graphical user interface (GUI, in MATLAB) for flexible and intuitive hardware control (Figure 1B and S2). To achieve reliable sampling at high frequencies with live display and without signal losses, we streamed the information of each photon, encoded in different bits of a 32-bit binary number, from the TCSPC boards to the computer. Individual photons were then assigned to individual lifetime bins (25–100 ps/bin) within a macroscopic sampling duration (0.1–5000 Hz sampling frequency). The total intensity, the mean photon emission time (MPET), and the lifetime decay curve (Figure 1C) can be displayed in real-time during recording. This integrated hardware and software system offers the following features: i) measurement at an intensity up to 2×10^7^ photons/s and a sampling rate up to 5000 Hz, ii) acquisition triggering by an external Transistor-Transistor Logic (TTL) pulse to enable synchronization with other experimental setups, iii) continuous acquisition durations up to 100 hours, iv) automatic recording loops at programmable time intervals, and v) integrated data visualization and data analyses.

### iFLiP accurately measures the lifetime of fluorophores at kHz sampling frequency

We first characterized the basic properties of the iFLiP system, including the instrument response function (IRF) and detector afterpulses, by measuring attenuated laser light fed directly into the detector (Figure S3). The IRF represents the combined timing spread from the laser pulse, detector, and electronics, and defines the temporal resolution of the system. Across intensities in multiple hardware systems, the IRF exhibited a full-width half-maximum (FWHM) of ∼0.21 ns (Figure S3A and S3B), corresponding to a sigma of 0.1 ns when approximated using a Gaussian distribution. This is an order of magnitude lower than the lifetime of most fluorescence sensors (∼1–3 ns), confirming the sufficient temporal precision of the system. Afterpulses, which are artifactual counts occurring at microsecond to millisecond timescales after a true photon arrival event in many types of detectors, contribute to an intensity-dependent uniform background in the laser repetition window (12.5 ns at 80 MHz). They were estimated to be 3–6% of the total intensity, specific to each hardware setup (Figure S3C). This afterpulse probability was largely consistent across the typical intensity range of iFLiP (Figure S3D), allowing it to be subtracted in analyses. In addition, other system background signals, which include a combination of light-independent dark counts of the detector, system autofluorescence, and Raman scattering from the fibers, could also be measured and subtracted from experimental data (Figure 2A).

**Figure 2.**
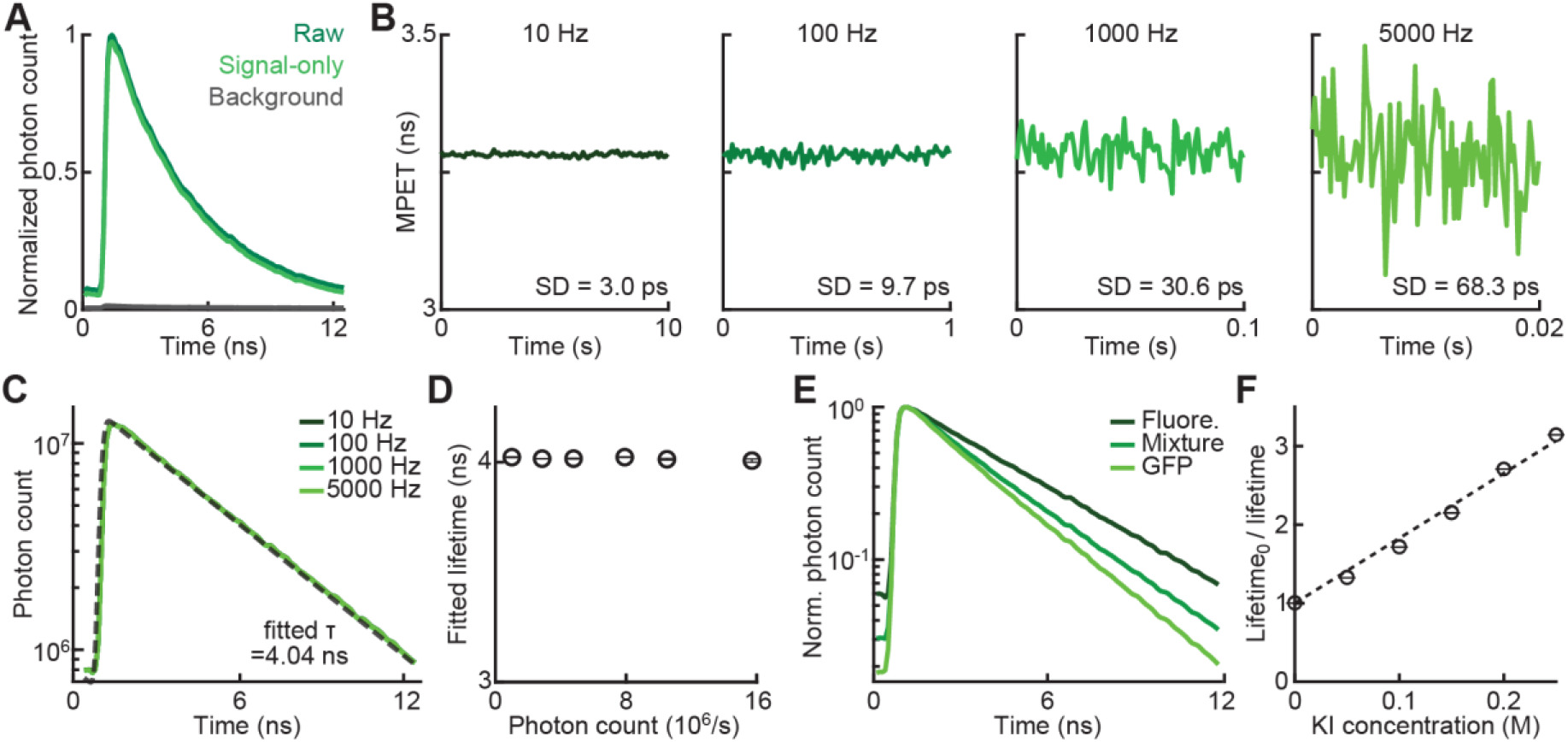
In vitro validation of iFLiP across sampling frequencies, intensities, and fluorophores. (A) Example normalized photon counts of raw fluorescein (0.1 µM), fluorescein after background subtraction (signal-only), and background. (B and C) MPET (B) and lifetime decay curve and the fit of the 10 Hz curve (dashed line) on the semi-log scale (C) of 5 µM fluorescein across sampling frequencies. (D) Fitted lifetime of fluorescein at various intensities resulting from different fluorescein concentrations (from left to right, n = 4, 4, 5, 8, 5, 5, respectively). (E) Normalized mean photon counts of fluorescein (0.5 µM, n = 5), GFP (0.5 µM, n = 3), and a mixture (n = 6) of fluorescein (0.125 µM) and GFP (0.375 µM). (F) Stern-Volmer plot of fluorescein (1 µM) quenching by KI (n = 3 for each concentration). Lifetime_0_ is measured in the absence of KI (n = 8). Dotted line: linear fit, lifetime_0_/lifetime = 8.278×[KI]+1. SD: standard deviation; KI: potassium iodide. Throughout, error bars: standard error. See also Figure S4.

To verify that iFLiP can reliably measure fluorescence lifetimes, we imaged well-characterized fluorophores in vitro (Figure 2). iFLiP was used to measure fluorescein across a broad range of sampling frequencies (10 to 5000 Hz). MPETs were consistent across these frequencies, although, as expected, fewer photons per time bin at higher frequencies resulted in increased standard deviations that matched theoretical predictions based on the photon counts (Figure 2B; STAR Methods). When integrated over the same duration, all sampling frequencies produced nearly identical lifetime decay curves. As expected from a single species of fluorophore, these curves were well fitted by a single exponential with a time constant (i.e., lifetime) of 4.04 ns (Figure 2C), consistent with previously reported fluorescein lifetimes in aqueous solutions.^27–29^ Furthermore, under the same excitation condition, the measured lifetime of fluorescein was consistent across four independent rigs (s.d. 0.038 ns, <1% of lifetime; Figure S4A) over a 15-fold change in the concentration and, therefore, in intensity (Figures 2D and S4B).

We then tested the analytical capability of iFLiP. First, we examined a mixture of two fluorophores: fluorescein and purified recombinant mEGFP. iFLiP measurements showed that the mixture exhibited a lifetime clearly distinct from that of each ingredient alone (fitted lifetime [ns; mean±sem]: fluorescein, 4.016±0.002; mEGFP, 2.699±0.005; mixture, 3.061±0.020) (Figures 2E and S4C). When fitting the mixture lifetime decay curve with two exponentials, the mEGFP-to-fluorescein fluorescence ratio was estimated to be 0.724:0.276. Based on the known extinction coefficient and quantum yield of these fluorophores (FPBase.org), a molar ratio of 3.05:1 was derived (STAR Methods), closely matching the mixture composition (0.375:0.125 μM). Additionally, in a separate experiment, iFLiP reliably detected a decrease in fluorescein lifetime upon potassium iodide (KI) quenching, which scaled linearly with KI concentrations to yield a Stern-Volmer constant of 8.3 (Figure 2F and S4D), in agreement with prior reports.^30,31^ Overall, these results indicate that the iFLiP system can reliably measure and analytically quantify the fluorescence lifetimes of multiple fluorophores.

### iFLiP reports fluorescence lifetime in biological samples

We next examined iFLiP’s ability to report fluorescence lifetime in biological samples using two widely employed preparations: cultured cells and acute brain slices. First, we expressed mEGFP or the cytosolic PKA lifetime sensor, tAKARβ^9^, in human embryonic kidney (HEK) 293 cells (Figure 3A and S5A). As expected, mEGFP expression resulted in a single exponential lifetime decay (Figure S5B) after subtracting background, which was measured when the fiber was pointed towards an area without expression. In contrast, the tAKARβ sensor exhibited multi-exponential lifetime decays (Figure 3B), as expected from its two activity states. Consistent with literature,^9,32^ iFLiP detected a decrease in the sensor lifetime (measured as MPET), indicating an increase in PKA activity, following the administration of the β-adrenergic agonist norepinephrine (1 µM), a physiological neuromodulator that activates the PKA pathway. The PKA response was further enhanced by strong activation of adenylate cyclases with forskolin (25 µM) and inhibition of phosphodiesterases with IBMX (50 µM) (Figures 3C and 3D).

**Figure 3.**
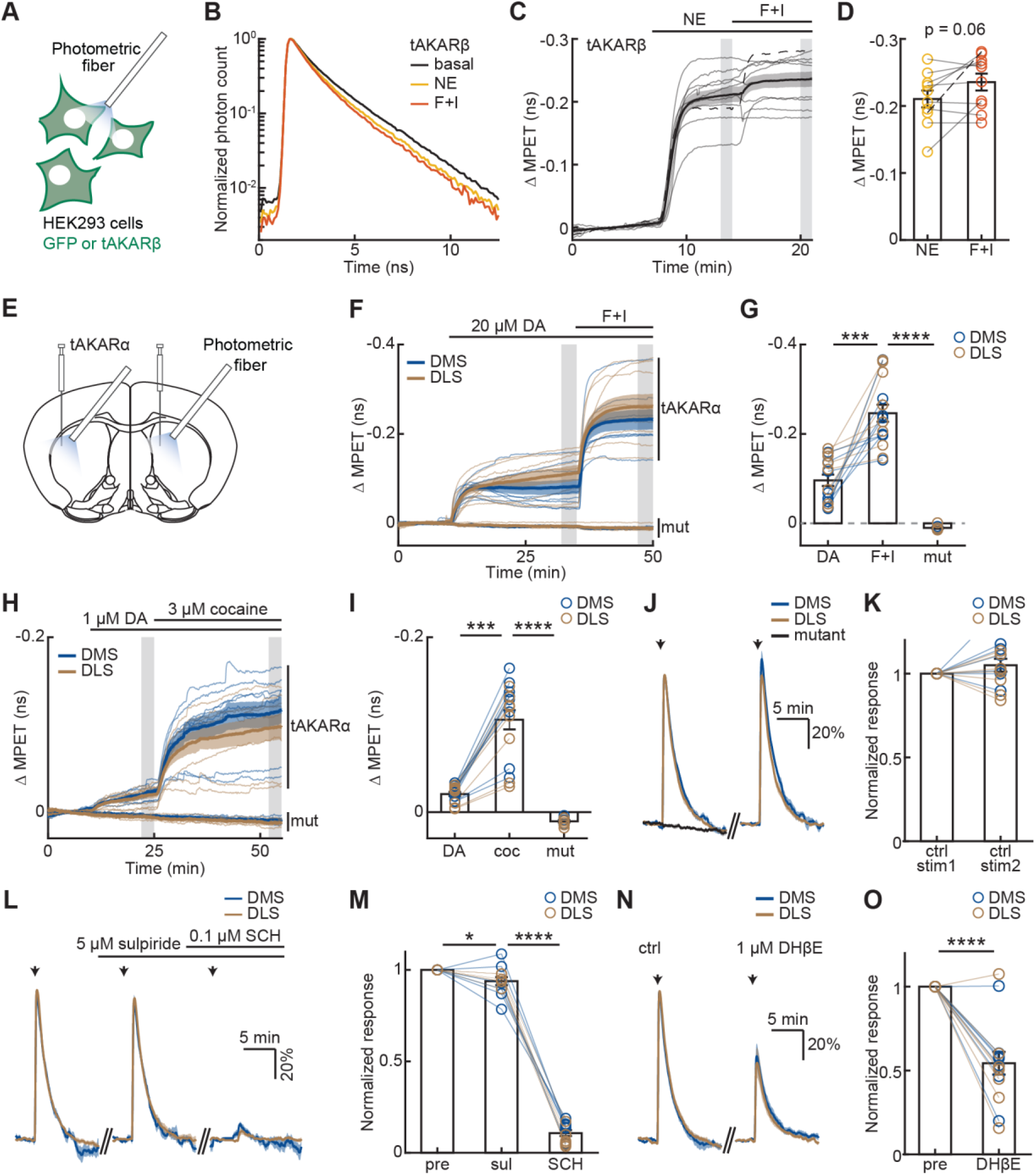
iFLiP monitors PKA responses in HEK cells and striatal slices. (A) Schematic of iFLiP recording in HEK cells. (B) Example normalized lifetime decay curve of tAKARβ in HEK cells under the indicated conditions, displayed on a semi-log scale. (C, D) Time course (C) and quantification (D) of ΔMPET of tAKARβ in HEK cells in response to the indicated drugs (n = 10 coverslips). Shaded rectangles: analysis time window for (B) and (D). Black dashed lines: the example recording analyzed for (B). (E) Schematic of iFLiP imaging in striatal slices. (F–I) ΔMPET of tAKARα or its mutant (mut) expressed in DMS (blue, tAKARα n = 8, mutant n = 4) and DLS (gold, tAKARα n = 8, mutant n = 5) slices in response to the indicated treatments. Gray shades in (F) and (H) indicate the quantification window for (G) and (I), respectively. mut: mutant tAKARα. (J–O) Electrically evoked (timing indicated by arrows) responses of tAKARα (colored) or its mutant (black, n = 5) in the DMS (blue) and DLS (gold) slices, without (panels J and K) or with the indicated drugs (L–O). n = 6 for (L) and (M), and 7 for other panels. F+I: forskolin (25 µM) + IBMX (50 µM); NE: norepinephrine (1 µM); DA: dopamine; coc: cocaine. Throughout, average (thick lines or bar), standard error (shadow or error bar), and individual data (thin lines or open circles) were shown. *: p < 0.05; ***: p < 0.001; ****: p < 0.0001. See Table S2 for detailed statistics. See also Figures S5 and S6.

Second, we used an adeno-associated virus (AAV) to express the microtubule-targeted PKA sensor, tAKARα^9^, in the dorsomedial and dorsolateral striatum (DMS and DLS, respectively), where PKA plays an essential role in mediating the neuromodulatory regulation of neuronal functions (Figure 3E).^33–37^ iFLiP measurements were performed in acute brain slices by placing an optical fiber tip towards the site of sensor expression (Figure S6A). The signal was readily detectable. Notably, when we simulated drifting by moving the fiber relative to the slice in the z dimension and simulated variable excitation conditions by changing the laser power, the intensity, but not lifetime readouts, were greatly affected in both cases (Figures S6B–S6E), demonstrating the advantages of iFLiP measurements.

### iFLiP reports physiologically relevant PKA activity dynamics ex vivo

Using acute striatal slice preparation, we asked whether iFLiP could detect dopamine-induced changes in PKA activity. Bath application of dopamine (20 µM) decreased the lifetime of tAKAR⍺ in both the DLS and DMS, corresponding to increased PKA activity. The PKA response was enhanced by forskolin (25 μM) and IBMX (50 μM) and, as a control, was absent when a phosphorylation-deficient mutant sensor was used (Figures 3F and 3G). In addition, inhibition of the dopamine transporter using cocaine (3 µM) enhanced the dopamine (1 µM)-dependent decrease in the lifetime of tAKARα (Figures 3H, 3I, S6F, and S6G).

Next, we asked whether iFLiP could detect PKA responses to endogenous dopamine release (Figures 3J–3M; see STAR Methods). Electrical stimulation elicited robust and reproducible lifetime responses using tAKARα but not the mutant sensor (Figures 3J and 3K). These responses were nearly abolished by the D1 dopamine receptor antagonist SCH23390 (1 µM) but not by the D2 antagonist sulpiride (5 µM) (Figures 3L and 3M), consistent with current knowledge of the different roles of D1 and D2 dopamine receptors in regulating PKA activity.^33–37^ Furthermore, the nicotinic acetylcholine receptor (nAChR) antagonist DHβE (1 µM) did not alter the basal PKA activity but significantly reduced these evoked PKA responses (Figures 3N, 3O, and S6H), supporting the recently appreciated role of nAChRs in mediating input-driven local dopamine release.^38–40^ Additionally, under our ex vivo conditions, there appeared to be no obvious differences in the PKA responses between the DMS and DLS.

Together, these results demonstrate that iFLiP reliably and stably reports sensor fluorescence lifetime changes in vitro and ex vivo.

### iFLiP reveals subregion– and cell-type-specific regulation of PKA in vivo

We next asked whether iFLiP could report fluorescence lifetime in vivo. We first expressed mEGFP in the DLS via AAV injection, and measured MPET at different sampling frequencies (Figure S7A). Similar to the in vitro measurements, iFLiP successfully measured the mEGFP lifetime at sampling rates up to 5000 Hz (Figures S7B–S7D).

We then evaluated whether iFLiP could quantify cell-type and region-specific PKA activity. Cre-dependent AAVs were injected into the DMS or DLS of *Drd1a-Cre* or *Adora2a-Cre* mice to express tAKARα in the direct and indirect pathway striatal spiny projection neurons, respectively (dSPNs and iSPNs, Figure 4A). iFLiP measurements were performed via implanted fiber cannulas under a head-fixed configuration (see STAR Methods). The sensor lifetimes were stable across measurement days (s.d. ∼6 ps; Figure S8) and were similar across the striatal subregions and SPN types, with the exception that DLS dSPNs had a slightly shorter lifetime (Figure 4B). In all conditions, the sensor lifetimes were significantly shorter than those of the corresponding mutant sensor (Figure 4B), suggesting that there are comparable levels of tonic PKA activity in both neuronal types. Furthermore, basal PKA activities across all conditions were correlated across weeks (Figure 4C), highlighting the stability of iFLiP measurements. Notably, quantification of these properties would be challenging with intensity measurements.

**Figure 4.**
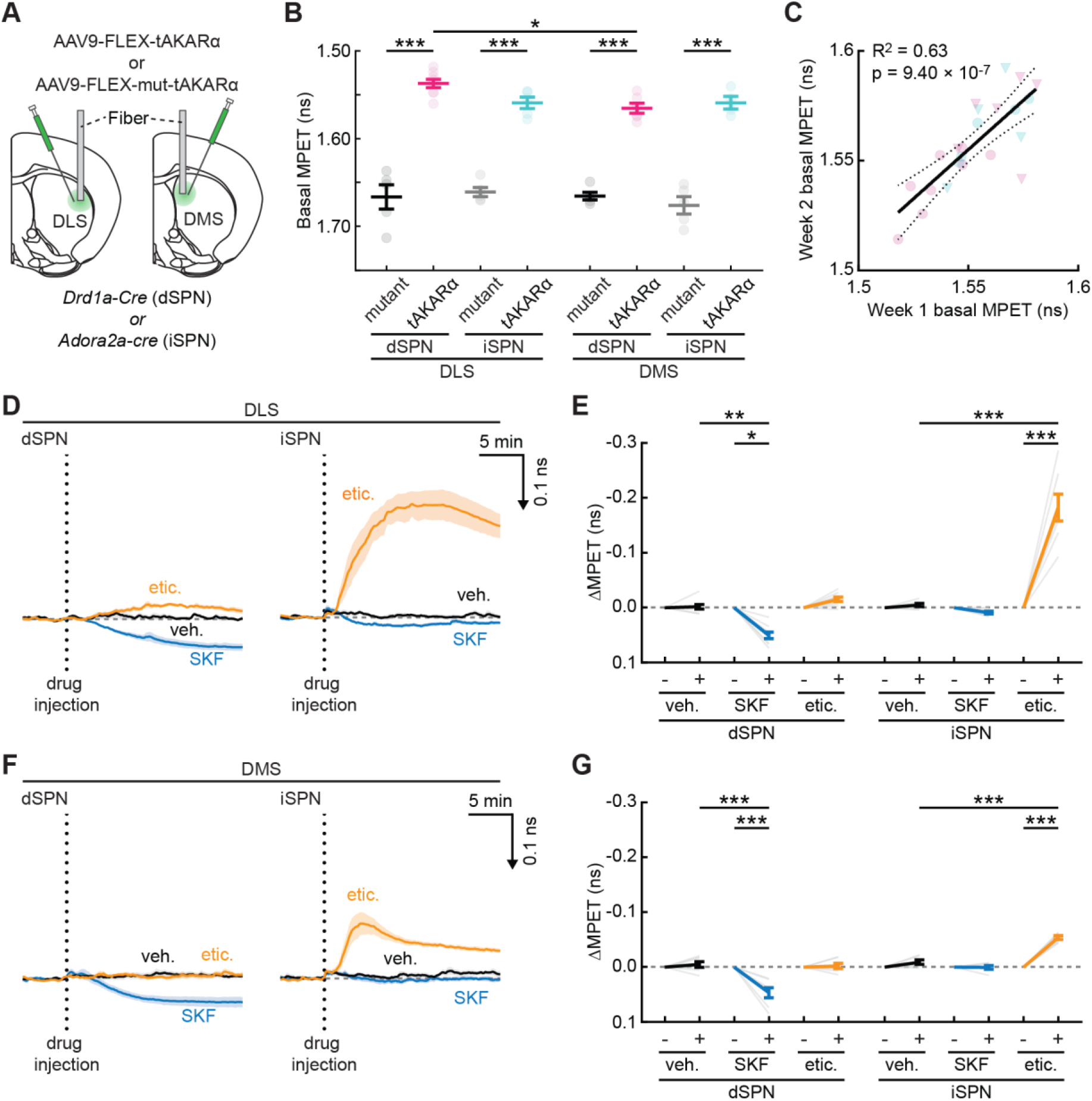
In vivo iFLiP measurements detect subregion– and cell-type-specific regulation of PKA activities in the striatum. (A) Schematics of virus injections and fiber cannula implantation. (B) Quantification of basal MPET of tAKARα or mutant tAKARα in dSPN or iSPN in the DLS or DMS. From left to right, n = 5, 8, 5, 7, 5, 6, 5, 6. (C) Correlation of basal MPET with linear fit (solid line) and 95% confidence intervals (dashed lines) of dSPN (magenta circle: n = 8) and iSPN (cyan circle: n = 7) in the DLS and dSPN (magenta triangle: n = 6) and iSPN (cyan triangle: n = 6) in the DMS across approximately a week. (D – G) Average traces and collective quantifications of basal MPET change in dSPN and iSPN in the DLS and the DMS by the indicated drugs. ΔMPET was calculated from the average of 15 to 20 min after drug injection. n = 9, 7, 7 and 6 for DLS dSPN, DLS iSPN, DMS dSPN and DMS iSPN, respectively. Veh.: vehicle; SKF: SKF83566; etic.: eticlopride. Throughout the figure, mean (darker lines) and s.e.m. (shaded area and error bars) are shown. *: p < 0.05, **: p < 0.01, ***: p < 0.001. See Table S2 for detailed statistics. See also Figure S7.

We examined how the observed PKA activities were modulated by dopamine receptors. It is known that D1 and D2 dopamine receptors are selectively expressed in dSPNs and iSPNs, respectively^35,37,41^. In both the DLS and DMS, systemic administration of the D1 antagonist SKF83566 (i.p.; 0.2 mg/kg) significantly reduced PKA activity, as reflected by increased sensor lifetime, without significantly affecting iSPN PKA activities (Figures 4D–4G). In contrast, the D2 receptor antagonist eticlopride (0.1 mg/kg) greatly enhanced iSPN PKA activity without significant effects in dSPNs. Notably, the effect of eticlopride on iSPNs was greater in the DLS than the DMS (Figure 4D–4G). Thus, different SPN subtypes exhibited distinct pharmacological sensitivities. Overall, iFLiP successfully monitored the dynamics of PKA sensor lifetime in vivo. These observations are consistent with the known roles of these receptors and cell types but also suggest previously unappreciated regional heterogeneity.

### iFLiP detects behavior-elicited PKA dynamics in vivo

We asked whether iFLiP could detect PKA dynamics induced by animal behavior using a head-fixed enforced locomotion paradigm (Figure 5A).^9,42,43^ In the DLS and DMS, the PKA activity in both dSPNs and iSPNs increased during enforced running, consistent with previous reports (Figure 5B–5D).^11^ Notably, dSPNs in the DLS exhibited an additional increase in PKA activity at the end of enforced running (Figures 5C, 5D, S9A, and S9B). This response was not evident in DMS, suggesting regional heterogeneity in locomotion-induced PKA dynamics. These lifetime responses were absent when a mutant PKA sensor was used (Figures 5C–5E). In both the DLS and DMS, locomotion-elicited lifetime responses were, on average, stable across weeks, although dSPNs in the DLS exhibited a small but statistically significant decrease (Figures 5F and S9A). Quantification of the kinetics revealed cell-type-specific differences. In both the DLS and DMS, the onset response of iSPNs (τ = 13 and 24 s, respectively) was significantly faster than that of the dSPN (τ = 32 and 38 s, respectively; Figures 5G and 5H). In contrast, the off-kinetics were comparable between dSPNs and iSPNs in both the DLS and DMS (Figures 5I and 5J).

**Figure 5.**
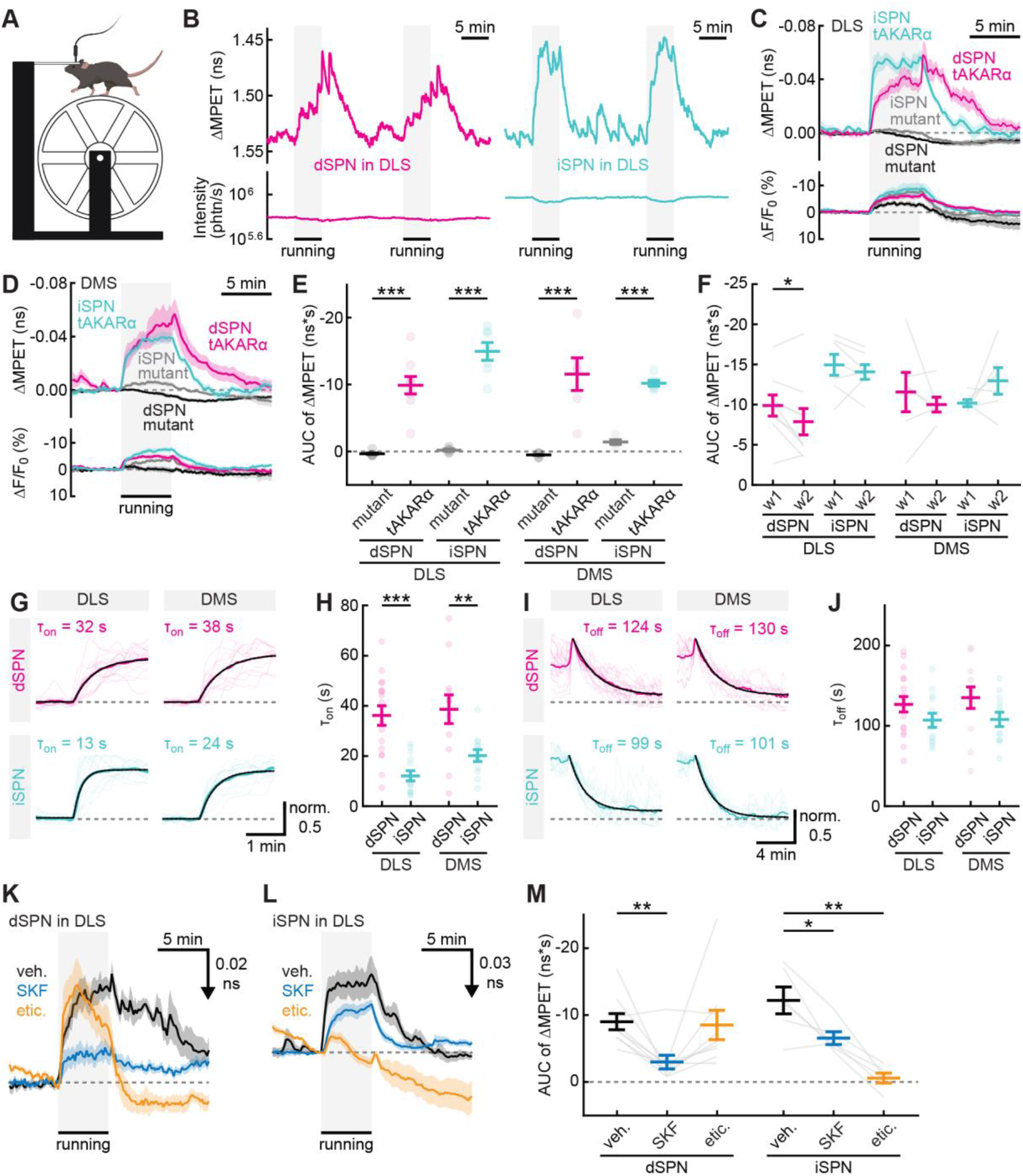
iFLiP detects locomotion-induced subregion– and cell-type-specific PKA dynamics in the DLS and DMS. (A) Schematic of mouse enforced wheel running during striatal PKA activity recordings with iFLiP. Created in BioRender. Yahiro, T. (2026) https://BioRender.com/u89hqpv. (B) Representative traces of tAKARα MPET and intensity changes in dSPNs (red) and iSPNs (cyan) of the DLS with enforced running. (C, D) Averaged traces of enforced running-induced MPET and intensity changes of tAKARα in dSPNs (C. red) and iSPNs (C. cyan) or mutant tAKARα (C. black) of the DLS, and tAKARα in dSPNs (D. magenta) and iSPNs (D. blue) or mutant tAKARα (D. gray) of the DMS. n = 8, 7, 6, and 6 for DLS dSPNs, DLS iSPNs, DMS dSPNs, and DMS iSPNs, respectively. (E) Quantification of MPET change of tAKARα and mutant tAKARα. n = 9, 7, 6, and 6 for DLS dSPNs, DLS iSPNs, DMS dSPNs, and DMS iSPNs, respectively. (F) Comparison of enforced running-induced MPET changes between week 1 (w1) and week 2 (w2). n = 9, 7, 6, and 6 for DLS dSPNs, DLS iSPNs, DMS dSPNs, and DMS iSPNs, respectively. (G – J) On-(G) and off-kinetic (I) traces of tAKARα in dSPN or iSPN in the DLS or DMS, and their quantification (H, J), including individual traces (light-colored), averaged traces (thick-colored), fitted curves (black solid traces), and the 95% confidence intervals (black dashed lines). For panels H and J from left to right, n (bout/mouse) = 17/9, 14/7, 12/6, 12/6, 17/9, 13/7, 12/6, and 12/6. (K–M) Pharmacological effects on enforced running-induced PKA changes of dSPN in the DLS (K) and iSPN in the DLS and DMS (L), and quantification (M). n = 9, 7, 6, and 6 for DLS dSPNs, DLS iSPNs, DMS dSPNs, and DMS iSPNs, respectively. W1: week 1; w2: week 2; veh.: vehicle; SKF: SKF83566; etic.: eticlopride. Throughout the figure, mean (darker lines) and s.e.m. (shaded area and error bars) are shown. *: p < 0.05, **: p < 0.01, ***: p < 0.001. See Table S2 for detailed statistics. See also Figures S8 and S9.

To examine whether dopamine receptor signaling contributes to locomotion-induced PKA activation, we systemically administered antagonists against D1 and D2 dopamine receptors while measuring tAKARα lifetime in DLS SPNs. As expected, mice in different experimental groups showed comparable responses to enforced locomotion before drug administration (Figure S9C). The D1 receptor antagonist SKF83566 greatly attenuated the running-elicited responses in dSPNs and, to a lesser degree, in iSPNs (Figures 5K–5M). In contrast, the D2 receptor antagonist eticlopride largely abolished the locomotion-elicited PKA responses in iSPNs but had only a moderate effect on those in dSPNs (Figure 4E). Given that the D2 receptor is Gi-coupled, this result was likely because eticlopride substantially elevated the basal PKA activity in these neurons (Figure S9D) to the degree that the sensor’s dynamic range was saturated. Overall, these results indicate that iFLiP enables robust lifetime measurements in behaving animals.

### iFLiP detects circadian PKA activity dynamics in the basal forebrain

We next examined whether iFLiP could report lifetime signals under more challenging experimental conditions, such as continuous, long-term recordings from deeper brain regions in freely moving animals. tAKARα was expressed in the basal forebrain via AAV injection (Figure 6A). Fluorescence intensity, fluorescence lifetime, and mouse motion were recorded simultaneously and continuously for 48 hours while allowing the animal to move or rest freely in a chamber (Figure 6B). The lifetime measurements were stable throughout the recording period, and tAKARα exhibited a lifetime significantly shorter than that of the mutant sensor (Figure 6C), reflecting a tonic level of PKA activity in these neurons. On average, the PKA activity was higher during the dark phase, when the animal was awake and active, than during the light phase. The lifetime of the mutant sensor also fluctuated, but to a lesser degree (Figures 6D, S10A, and S10B), likely due to the fluorescent protein’s sensitivity to internal temperature^44,45^, which may fluctuate across sleep-wake cycles. At a finer temporal resolution, tAKARα detected a nearly bimodal distribution of PKA activity, with higher activity closely correlating with animal movement and lower activity matching animal quiescence (Figure 6B). Notably, these lifetime responses exhibited a much higher signal-to-noise ratio than the corresponding intensity measurements (Figure 6E).

**Figure 6.**
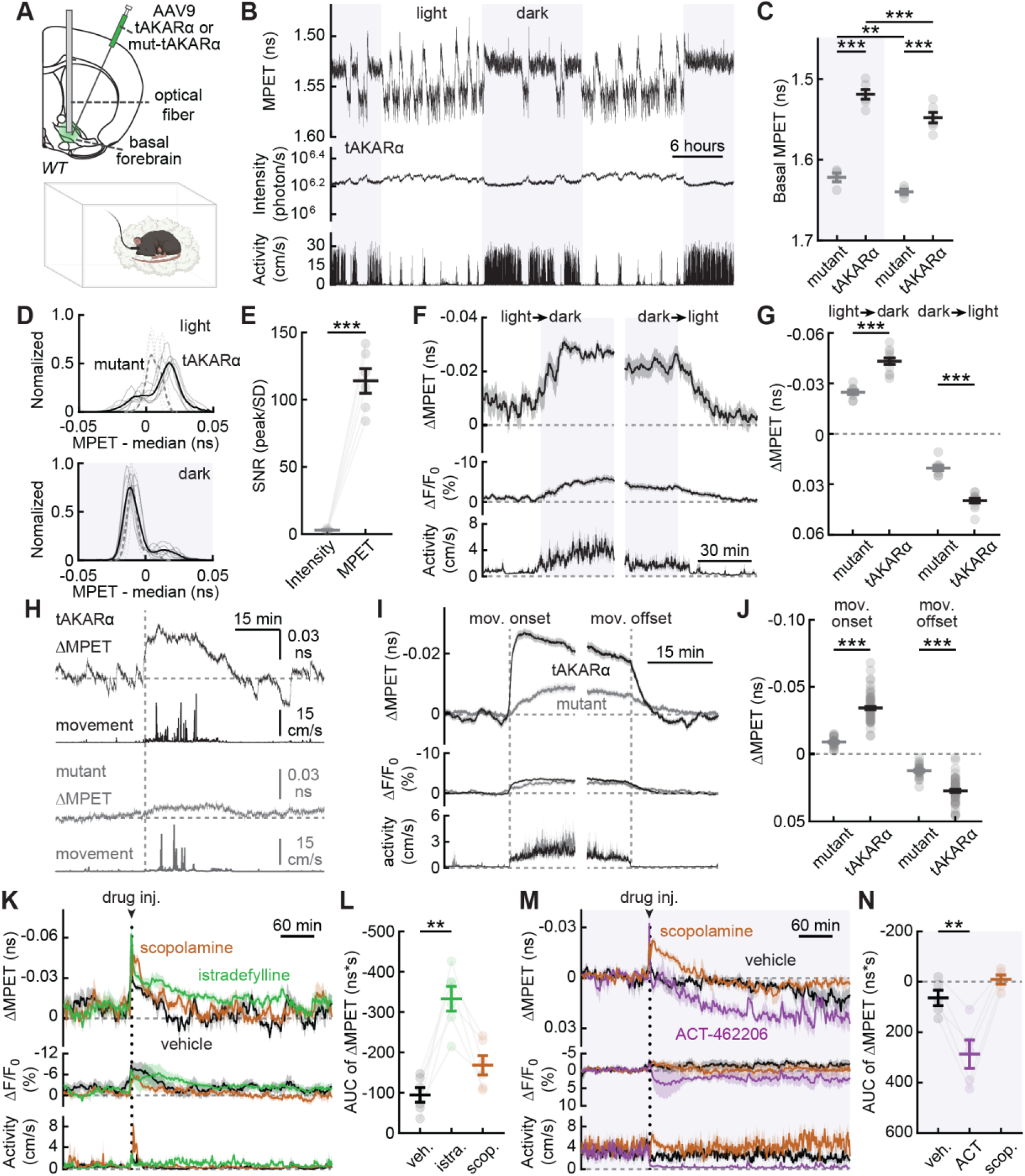
Longitudinal iFLiP recording detected circadian rhythmic PKA activity in the basal forebrain. (A) Schematic of virus injections and fiber cannula implantation into the basal forebrain (top) and longitudinal recordings to assess PKA activity across sleep-wake cycles (bottom; Created in BioRender. Yahiro, T. (2026) https://BioRender.com/ue2zin1). (B) Representative traces of a 48-hour recording of tAKARα in the basal forebrain. (C) Basal MPET shifts between the light and dark (without and with gray background, respectively) phases. n = 4 and 6 for mutant and tAKARα, respectively. (D) MPET distribution of mutant (dashed line) and tAKARα (solid line) during the light phase and dark phase. Individual traces are also shown (light-colored lines). (E) Comparisons of signal-to-noise ratio between intensity and MPET during the light phase. n = 6. (F) Averaged traces of tAKARα MPET, intensity and locomotor activity changes during light-to-dark and dark-to-light phase transitions. (G) Quantification of MPET and intensity changes during the light-to-dark and dark-to-light phase transitions. n = 4 and 6 for mutant and tAKARα, respectively. (H) Representative traces of spontaneous PKA activity accompanied by locomotor activity during the light phase. (I) Averaged traces of spontaneous PKA activity during the light phase aligned with the onset (left) and offset (right) of the locomotor activity. (J) Quantification of the MPET change at the onset and offset of the locomotor activity. n (bout/mouse) = 25/4, 66/6, 26/4 and 62/6 from left to right. (K-N) Time course traces (K and M) of averaged ΔMPET and ΔF/F0 of tAKARα in the basal forebrain aligned to the locomotor activity and the AUC quantification of ΔMPET (L and N) after i.p. injections of the indicated drugs during the light (K and L) or dark (M and N) phase. n = 6 for K and L; n = 5 for M and N. Veh.: vehicle; istra.: intradefylline; scop.: scopolamine: ACT: ACT – 462206. Throughout the figure, mean (darker lines) and s.e.m. (shaded area and error bars) are shown. *: p < 0.05, **: p < 0.01, ***: p < 0.001. See Table S2 for detailed statistics. For panels D, F, I, n = 4 and 6 for mutant and tAKARα, respectively. See also Figures S10.

At the light–dark transitions, tAKARα detected increased PKA activity at the light-to-dark transition and a decrease in PKA activity at the dark-to-light transition (Figures 6F and 6G), presumably reflecting changes in the animal’s internal states. In addition, during the light phase, tAKARα detected transient increases in PKA activity when the animal exhibited short bouts of intermittent locomotor activity (Figure 6B and 6H–6J).

Finally, because an animal’s light-dark cycle transitions are associated with its circadian rhythms, we pharmacologically manipulated the sleep–wake signaling while monitoring basal forebrain PKA activity. First, we examined the effects of istradefylline, an A_2A_ receptor antagonist known to disrupt sleep, during the light phase when mice sleep more. Systemic istradefylline administration (5 mg/kg) induced a long-lasting, significant increase in basal forebrain PKA activity accompanied by increased locomotor activity (Figures 6K, 6L, and S10C). In contrast, scopolamine, a muscarinic receptor antagonist, only transiently altered PKA activity and locomotion under the same conditions (Figures S10D and S10E), presumably due to different pharmacokinetics or circuit-level mechanisms. We next tested ACT-462206, a sleep-promoting orexin receptor antagonist, during the dark phase when the animal is typically active. ACT-462206 reduced both PKA activity and locomotor activity (Figures 6M, 6N, and S10F). In contrast, scopolamine only elicited transient effects (Figures S10G and S10H). Together, these results demonstrate that iFLiP enables continuous monitoring of PKA dynamics in deep brain regions and can detect both circadian and pharmacologically induced changes in neuromodulatory signaling in freely moving animals.

## DISCUSSION

Here, we present a simple, low-cost, yet flexible, robust, and turn-key system for lifetime fiber photometry. Although the advantages of fluorescence lifetime measurements have long been recognized, and lifetime measurements have been implemented with microscopy and fiber photometry in certain laboratories,^10,15,19^ this technology remains inaccessible for most laboratories because it requires complex, expensive custom setups and specialized expertise in optics and electronics. iFLiP addresses this critical need by providing a simple hardware design and user-friendly software for turn-key operation. The total cost (∼$22k, Supplemental Table 1) is over 40% lower than that of any previously published systems.^46^ At the same time, this system achieves robust performance, including a sampling rate of up to 5000 Hz and a maximum photon count of 2×10^7^/s. At 20 Hz, this corresponds to up to 1×10^6^ photons per sample point and 0.1% precision in lifetime quantification (*δτ*/*τ* = 1/√*N*; STAR Methods). Given that many modern sensors exhibit lifetime dynamics of 25% or more,^9,12,13,17,47^ the sensitivity of the current system enables reliable lifetime detection across a broad range of applications. In addition, the system has an IRF with a FWHM well below the lifetime of most sensors and fluorescent proteins. This is essential as the IRF defines the lower limit of detectable lifetime, analogous to how the point spread function defines the resolution of microscopes.

The system presented here is designed for single-channel detection of green fluorescence (e.g., EGFP). The laser and filters can be adjusted to detect sensors of other colors. Multiplexed imaging of multiple sensors or simultaneous data collection at multiple brain regions using a multi-channel TCSPC board is, in principle, possible. However, this is often associated with greatly increased cost and light path complexity. Future efforts to simplify the design for robust multiplexed imaging will be needed.

Compared to lifetime measurements based on two-photon microscopy, fiber photometry exhibits significantly higher background signals. As the excitation light is required to pass through meter-long optical fibers, fiber autofluorescence and Raman scattering become substantial. Careful selection of the emission filter is therefore critical. The dark count of the detector is also non-negligible. Another source of background is the detector afterpulses, which scale with the detected intensity. Other detectors, such as photomultipliers (PMTs) or single-photon avalanche diode (SPAD), have lower afterpulse probabilities. However, these alternatives add significant cost and have their own limitations: PMTs may become saturated or damaged at high intensities, whereas SPAD detectors often have a very small active area, making light collection challenging. In iFLiP, hardware optimization combined with background and afterpulse subtraction still enables accurate lifetime measurements (e.g., Figure 2).

When compared to systems based on two-photon microscopy, the iFLiP approach is simpler, more broadly accessible, more readily compatible with freely moving paradigms, and less invasive for imaging deep brain regions (ϕ = 200 μm for optical fibers, c.f. 1400 μm for gradient-index [GRIN] lenses with a cannula), although the gains come at the expense of single-cell resolution. By using iFLiP to measure PKA activity in the striatum, we obtained several observations that extended the current understanding of striatal neuromodulation. First, in acute brain slices, PKA activity in the DMS and DLS exhibited similar sensitivity to dopamine and cocaine manipulations ex vivo. Thus, the recently reported spatial heterogeneity in dopamine release across striatal regions likely propagates to intracellular signaling activity.^48,49^ Second, our result that an nAChR antagonist attenuated PKA activation induced by electrically elicited dopamine release corroborates recent observations that nAChRs expressed on dopaminergic axons mediate local dopamine release driven by cortical inputs.^38–40^ Additionally, in vivo iFLiP measurements in the striatum largely verified our previous observations using in vivo two-photon fluorescence lifetime imaging that animal locomotion drives cell type specific PKA signaling.^11^ Furthermore, this approach allowed us to readily compare responses across multiple striatal subregions. We found that PKA activity of dSPNs in the DLS, but not the DMS, exhibited a clear peak at the offset of enforced animal locomotion, which may be related to recently described dopamine release that occurs upon stress relief.^50^ Finally, we found that PKA activity on-kinetics in iSPNs were much faster than those in dSPNs. Such differential kinetics may provide a dynamic temporal window critical for striatal function, such as movement control, warranting future studies.

The iFLiP system measures fluorescence lifetime by counting individual photons with high sensitivity (quantum efficiency ∼40%). Because of this sensitivity, in vivo iFLiP experiments often require excitation powers an order of magnitude lower than those of typical intensity-based fiber photometry experiments (e.g., less than 1 μW was used for all experiments in Figure 6). This, together with the aforementioned advantage of lifetime measurements, allows for long-term continuous recordings in freely moving mice. Such recordings in the basal forebrain revealed PKA activity dynamics correlated with the animal’s circadian rhythm: tonic PKA activity increases in the dark phase but decreases in the light phase. The potential role of this PKA activity in driving sleep-wake state transitions is supported by its sensitivity to manipulations of arousal-related neuromodulatory pathways, including the adenosinergic, cholinergic, and orexinergic systems. Future experiments combining iFLiP with EEG/EMG, cell-type-specific recordings, and receptor-specific manipulations will be necessary to define the receptor, cellular and circuit mechanisms by which PKA bridges neuromodulatory activation and animal behavior. Nevertheless, our results demonstrate that iFLiP can reveal previously unresolved intracellular signaling states in freely moving animals that are important for brain function.

### Limitations

Multiple factors collectively impose a practical limit on the maximal intensity measurable with iFLiP. First, although the system minimizes the distortion by TCSPC deadtime in lifetime determination by implementing a compatible laser frequency, photon losses caused by the deadtime still limit the collectible light flux. Second, the detector has a finite speed, such that each photon generates a pulse with a FWHM of ∼4 ns (Thorlabs.com). Individual photons arriving within this time duration cannot be resolved from each other, thereby effectively imposing another deadtime that may distort lifetime at high signal intensities. Third, the need to process every photon also imposes challenges for data transfer and CPU computation at high count rates. Achieving a higher intensity limit using the TCSPC mechanism will require improvements on all of these fronts. Nevertheless, as demonstrated, the current intensity limit is sufficient for most in vivo lifetime measurements.

Similar to other single-photon excitation methods, the autofluorescence from brain tissue can be substantial. Although the background from the hardware system is typically stable and can be measured and subtracted, background from the sample, particularly in vivo, is usually variable across experiments and can be challenging to remove reliably. Additionally, although lifetime measurements are more resistant to factors affecting fluorescence intensity, they might still be perturbed by certain biological factors. For example, it is known that temperature may affect the lifetime of many fluorophores^44,45^. Therefore, iFLiP results should be compared with those obtained from a corresponding functionally deficient mutant sensor and interpreted with care.

## SUPPLEMENTAL INFORMATION

Supplemental Information includes ten figures and a supplementary table.

## AUTHOR CONTRIBUTIONS

H.Z. conceived the project and developed the initial setup. T.Y., Y.C., L.B.E., T.M., and H.Z. designed the experiments. T.Y., Y.C., L.B.E., S.A., and H.Z. performed the experiments and analyzed the data. T.Y., Y.C., L.B.E., and H.Z. wrote the original draft of the manuscript. All authors revised and edited the manuscript. T.M. and H.Z. supervised the project and secured funding.

## ACKNOWLEDGEMENTS

We thank Zilong Liu and Qing Qu for associated efforts that are not included in the manuscript, Drs. Stephan Lammel, Garret Stuber, and Charles Zhou for critical comments on the manuscript, Drs. Isabelle Baconguis and John Williams for sharing fluorescent samples, Drs. Robert Spencer and Ali Mohebi for feedback on the iFLiP software, Maozhen Qin for mouse husbandry, and all members of the Mao laboratory and Zhong laboratory for helpful discussions. This work originated from a teaching project at the *Neurobiology* summer course of Marine Biological Laboratory, Woods Hole, Massachusetts, and was supported by three NIH BRAIN Initiative awards UM1MH138931(H.Z. and T.M.), RF1NS133599 (H.Z. and T.M.), RF1MH130784 (H.Z.), and two NINDS R01 grant R01NS081071 (T.M.) and R01NS127013 (H.Z.).

## STAR METHODS

### Contact for reagent and resource sharing

Further information and requests for resources and reagents should be directed to and will be fulfilled by the Lead Contacts, Tianyi Mao and Haining Zhong.

### iFLiP system

The minimum essential components for iFLiP and their sources are listed in Supplementary Table 1. The optical components are connected to each other using standard optical fibers (diameter 200 μm, numeric aperture [NA] 0.37; also listed in the table) via FC/PC connectors as shown in Figure 1A. The detector output and laser sync signals were connected to the appropriate channels on the TCSPC board via coaxial cables with SMA connectors. The iFLiP software (current version iFLiP2), written in MATLAB 2022a and compiled as a standalone program, will be made available on GitHub.

### Estimation of the impulse response function (IRF) and afterpulse probability

To estimate the IRF and afterpulse probability, we fed attenuated (∼7 optical densities) laser light directly to the detector. The IRF was visualized as the summed photon curve within the laser repetition window after dark-count subtraction (Figure S3). A Gaussian distribution was fitted to the IRF to calculate the standard deviation. The same photon curve was used to calculate the afterpulse probability: the contribution of afterpulse counts was estimated as the count at the lowest 2% across all time bins multiplied by the total number of time bins. The threshold is subjectively selected to eliminate the effect of incoming signal and autofluorescence. Alternatively, the afterpulse probability can be estimated as a constant component in fitting the lifetime curve of a fluorophore with a known decay time after subtracting other backgrounds. Both approaches give comparable results (difference <1% in the afterpulse probability value).

### Lifetime correction and analysis

Unless noted otherwise, all observed intensity data (*N_obs_*) were corrected for afterpulses (*N_ap_*) and background (*N_bg_*) before further analysis:

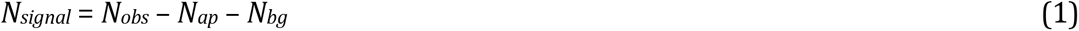

Photon counts resulting from afterpulses (*N_ap_*) were assumed to be proportional to the macroscopic light intensity (*R_obs_*):

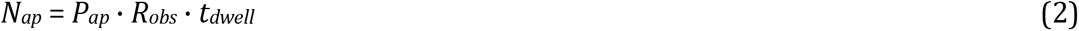

Where *P_ap_* is the afterpulse probability and *t_dwell_* is the dwell time of the particular time bin for *N_ap_*.

To estimate *N_bg_*, we measured the apparent signal in the absence of the sample. For in vitro experiments, this entailed measuring the background optics including the fiber used to capture the signal from the cells in the bath. For ex vivo experiments, this entailed measuring the background by pointing the fiber toward a slice area without expression. For in vivo experiments, this entailed measuring the background of the optics, plus an implantable fiber cannula. Because signal intensity is much larger than the background (usually 1–2 orders of magnitude), the fraction of background in the signal is lower than the background measured at low photon counts because of the loss of photons due to TCSPC deadtime (*τ_dead_*). The light collection efficiency is given by

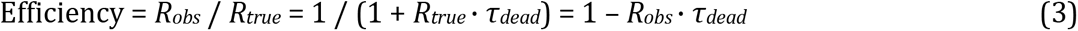

Where *R_true_* is the true incident photon rate. Since the efficiency is high for most background-only measurements (typically <10^6^ photons/s; efficiency > 97%), we use the measured background (*N_bg, measured_*) as it is, without correction, to calculate the effective background under a given signal intensity:

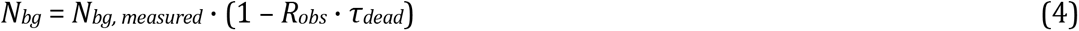

Combining Equations (1), (2), and (4), the true fluorescent signal can be estimated as:

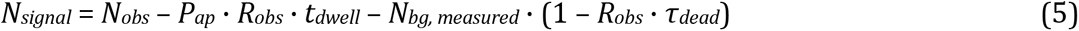

Lifetime was measured from the background-subtracted, afterpulse-corrected intensity data. Two different measurements are used: the time constants (i.e., fitted lifetimes) and mean photon emission time (MPET), with the latter being more computationally efficient and more repeatable (e.g., insensitive to non-convergent fits). Fitted lifetimes were determined by fitting single or double exponentials to the photon count curve over time convoluted with the impulse response function (approximated as a Gaussian distribution) (Figures 2E and 2F).

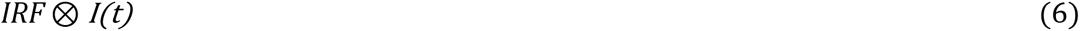

Where IRF is approximated using a Gaussian distribution:

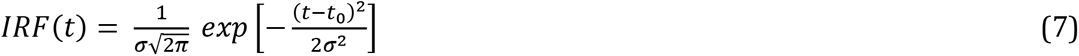

And I(t) is

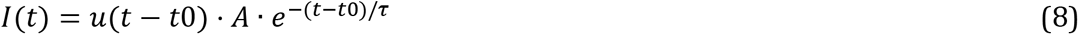

for a single exponential decay, or

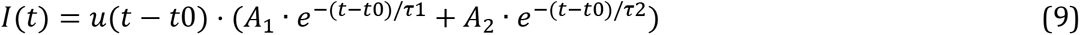

for a double exponential decay, where *u*(*t*) is the Heaviside step function. To correct for the incomplete lifetime decay from previous pulses, the effect resulting from previous pulses up to 10 times of the larger τ before the current pulse is considered in fitting.

MPET was calculated as

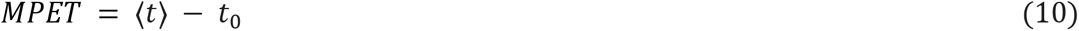

Where *t* is the arrival time of individual photons and *t*_0_ is the timing of the laser pulse (derived from fitting). Note that t0 can be adjusted by changing sync or signal offset and is typically set to ∼1 ns, so that the decay curve utilizes the majority of the repetition window (12.5 ns for an 80 MHz laser). For single exponential decays, MPET equals the fitted lifetime. For double exponential decays, MPET is related to the fitted lifetimes with the following equation:

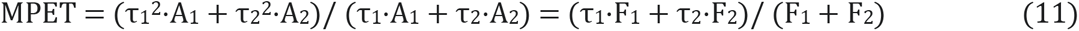

where F_1_ and F_2_ are the total intensities of the components with lifetime of τ_1_ and τ_2_, respectively.

Due to a finite integration window (t_w_; smaller than the repetition window, which is 12.5 ns for an 80 MHz laser), the apparent MPET is underestimated. For example, for a single exponential decay, MPET should be identical to fitted lifetime. However, the apparent MPET is smaller:

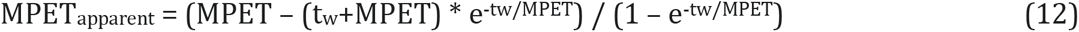

For mEGFP (lifetime ∼ 2.6 ns) and a typical measurement window of 11 ns, the apparent MPET is 2.44, or 94% of the real value. In this manuscript, all MPETs are MPET_apparent_ without correction.

For in vitro and ex vivo recordings, the SPC range used for MPET calculation was typically 0.4–12.3 ns. For in vivo recordings, the range was typically restricted to 0.4–10 ns to avoid a minor background peak beyond 10 ns, likely arising from reflection of excitation light and thereby their timing at surface junctions (e.g., the rotary joint connection). The SPC range duration was matched for MPET comparisons.

The noise level of lifetime determination was predicted to be:

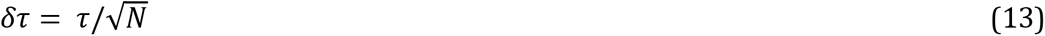

Where N is the number of detected photons^7,9^.

For analysis of the mEGFP-fluorescein ratio, the data were fitted by two exponentials (Equation 9). The total fluorescence of each component is given by *A* · *τ*. The extinction coefficient (at max excitation wavelength), the relative excitation strength (at 486 nm, peak spectrum under our application settings; Thorlabs.com), and emission quantum yield of each fluorophore were 56,000 M⁻¹cm⁻¹, 0.99, 0.60 for mEGFP and 88,000 M⁻¹cm⁻¹, 0.71, 0.92 for fluorescein (FPbase.org). This gives a relative brightness of 1:1.73 (mEGFP:fluorescein).

### Cell culture and transfection

HEK-293 cells were obtained from American Type Culture Collection (ATCC #CRL-1573). Note that all cells from ATCC have been authenticated by morphology, karyotyping, and PCR-based approaches and tested for mycoplasma. Cell aliquots (within 2–4 passages) were kept frozen in liquid nitrogen until use. Thawed cells were further authenticated within the lab based on their morphology and growth characteristics and were maintained in 100-mm cell culture dishes (Fisher Scientific, #FB012924) at 37 °C with 5% CO2 in minimal essential medium (ThermoFisher #11095-080) with the addition of 10% fetal bovine serum. Each aliquot of cells was passed and used for no more than 4 months.

To record from transfected HEK cells under the iFLiP setup, cells were subcultured onto glass coverslips (∼ 5 x 5 mm, hand-cut using a scriber) coated with 0.1 mg/mL Poly-D-Lysine (Sigma, #27964-99-4) in 35-mm cell culture dishes (Corning, #CLS430165), or seeded in 4-Chamber 35-mm glass bottom dishes (Cellvis, #D35C4-20-1.5-N). For sensor expression, tAKARβ in mammalian expression plasmids (0.3 µg/35-mm dish) was transiently transfected using Lipofectamine-2000 (ThermoFisher, #11668030) according to the manufacturer’s instructions with the exception that only 3 μl of the reagent was used and measured at two days post-transfection.

### In vitro photometric recordings

To record from fluorophores in vitro, a Doric patch cable (200 µm, NA 0.37, low autofluorescence) was cut near the distal end and stripped to expose ∼2 cm of the inner glass fiber. Care was taken to ensure a relatively smooth cut surface, minimizing light scattering. Before recordings, the iFLiP hardware was allowed to warm up for 1–3 hours or longer, and the fiber was bleached until the autofluorescence reached a plateau. Depending on the setup and laser power, this took between 15 minutes (3 mW) and 90 minutes (40 µW). To measure background, the laser power was kept consistent while fluorescence was measured with the sample absent—either in solvent alone or in a bath without the cell coverslip present.

To record the lifetime of fluorescein and mEGFP, the exposed fiber end was submerged into fluorescent solutions (0.6–1.6 ml) of fluorescein, GFP, or a mixture of the two made in ddH_2_O (Figure 2B and 2C) or 25 mM HEPES-buffered HBSS solution (pH = 8, Figure 2A, 2D–2F) in a 4-chamber glass-bottom Petri dish (Cellvis). The laser power (typically 0.3 – 10 μW) was adjusted such that the measured photon counts were ∼0.1×10^7^ to 1×10^7^. To record from fluorophores expressed in HEK cells, the coverslip was placed on the bottom of the chamber perfused with carbogen (95% O2/5% CO2) gassed artificial cerebrospinal fluid (aCSF) containing (in mM) 127 NaCl, 25 NaHCO_3_, 25 D-glucose, 2.5 KCl, 1.25 NaH_2_PO_4_, 2 CaCl_2_, and 1 MgCl_2_. The chamber was covered in black electrical tape to reduce ambient light. The fiber was placed in an area with medium expression and over cells with healthy morphology. Specifically, the fiber tip was lowered until it touched the coverslip and was then retracted slightly. The measured photon counts read ∼3×10^6^ to 10×10^6^. Drugs were bath-applied. The fiber and chamber were washed between experiments with 70% ethanol and ddH_2_O.

### Experimental animals

All animal experiments were conducted according to the US National Institutes of Health guidelines for animal research and were approved by the Oregon Health & Science University Institutional Animal Care and Use Committee (IACUC protocols IP00000955 and IP00002274). The following mice were used in this study: wildtype C57BL/6 mice (Charles River Laboratories, Wilmington, MA, USA); *Drd1a-Cre* (Mutant Mouse Resource and Research Centers, 036916) and *Adora2a-Cre* (Mutant Mouse Resource Research Centers, 036158). All transgenic mice were backcrossed at least five generations to a C57BL/6 background. Mice of 1.5-8 months old with approximately equal proportions of both sexes were used. All mice were housed in groups when possible and maintained in a standard 12-hour light/dark cycle with ad libitum access to food and water. All in vivo experiments were performed in the light cycle, except for freely moving experiments, which were performed in both light and dark cycles. Mice were randomized to treatment groups, and sexes were approximately equal between groups.

### Stereotaxic viral injections

The following viruses were used: AAV9-CAG-FLEX-cd-tAKARα (custom production; 1.16×10^12^ GC/mL), AAV9-hSyn-cd-tAKARα (custom production; 1.98×10^12^ GC/mL). The flexed virus was diluted 1:5 in PBS prior to injection. 150-200 nL of the diluted viral vector was injected into one of the following locations using a glass pipette, as previously described^51^. The coordinates are (in mm relative to bregma): for the DMS, 0.4–0.7 anterior, 1.1–1.15 lateral, 2.8 ventral; for the DLS, 0.2–0.5 anterior, 2.4 lateral, 2.8 ventral; for the basal forebrain 0.5 anterior, 1.2 lateral, 5.0 ventral. Stereotaxic injection was performed under isoflurane anesthesia (5% induction, 2% maintenance). Mice were administered 5mg/kg carprofen for 3 days following surgery and were allowed to recover for 1-3 weeks prior to fiber implantation.

### Ex vivo slice preparation and recordings

Wildtype animals injected with codon-diversified (cd)-tAKARα in the DMS and DLS were sacrificed at P40-60 and decapitated for brain extraction. Brains were sliced coronally on a vibratome (VT1200S, Leica Biosystems, Nussloch, Germany) at 300 µm thickness in fresh ice-cold cutting solution (in mM: 25 NaHCO_3_, 1.25 NaH_2_PO_4_·H_2_O, 25 glucose, 2.5 KCl, 7 MgCl_2_·6H_2_O, 0.5 CaCl_2_, 110 choline chloride, 11.5 sodium ascorbate, and 3 sodium pyruvate). Slices were incubated at 37 °C for 30 minutes and then at room temperature for 0.5–10 hours in carbogenated aCSF before use.

Brain slices were separated along the midline to isolate two half slices each containing one hemisphere. A half slice was then stabilized to the perfusion chamber with a custom-made U-shaped harp. The recording fiber (at a 30-degree angle to the slice) was lowered into the bath toward the expression center, or an area without expression as a control, in the DMS or DLS, until positioned right above the slice. For the intensity drift experiments, the fiber was moved upward for 10 µm to induce intensity change and then moved back to baseline position (Figures S6B and S6C). Alternatively, with the fiber position unchanged, the baseline laser intensity (∼0.8–2µW) was decreased by 5–10% using an inline optical attenuator (V450A, Thorlabs; controlled by a voltage signal generator, TKXEC, QH-VISG2-ED) and then restored to baseline (Figures S4D and S4E). All drugs were bath-applied at the following concentrations (in µM): dopamine 20 or 1, forskolin 25, IBMX 50, cocaine 3. Dopamine and cocaine stocks were prepared in ddH_2_O, and forskolin and IBMX were dissolved in DMSO. The chamber and fiber were cleaned with at least 6 rounds of wash with ddH_2_O and 70% ethanol between experiments. The pharmacology experiments (Figures 3F–3J) were performed at room temperature.

For electrical stimulation experiments (Figures 3J–3O), a theta-tube was pulled from capillary glass tubing (TG150-4, Warner Instruments), connected to a stimulation box, and positioned below the slice surface 150–200 µm away from the fiber tip at 32–34 ℃. A train of 3 electrical stimulations (25–75 µA) was delivered at 25 Hz with a dwell time of 0.1–0.5 ms to induce an MPET change typically between 0.03 ns and 0.12 ns. A cocktail of glutamatergic and GABAergic blockers (in μM: 10 NBQX, 10 CPP, 20 SR95531, and 1 CGP55845) was added to the bath to isolate putative dopamine-induced responses. CGP55845 stock was prepared in DMSO; NBQX, SR95531, and CPP stocks were prepared in ddH_2_O. Parameters were kept consistent across stimulations within each experiment. For the mutant control, parameters matched the strongest stimulation applied to the experimental group on the same day. Drugs were applied at least 5 minutes after a stable baseline was reached and incubated for at least 15–20 minutes before the next stimulation.

### Fiber implantation

For in vivo photometry experiments, optical fiber cannulas were implanted under isoflurane anesthesia (5% induction, 2% maintenance). 1–2 weeks after stereotaxic viral injections, fiber cannulas (200 μm diameter, 3.5mm long, 0.39 NA; RWD) were implanted at the same coordinates as the injection site by inserting them while recording signals with a custom-built spectrally resolved photometry system^52^. Using a 488 mm LED to excite green fluorophores, the spectrum was observed during fiber implantation, and the fiber was implanted to the depth of the greatest signal and secured with dental cement mixed with cyanoacrylate. Mice were administered 5 mg/kg carprofen for three days following surgery and were allowed to recover for at least two weeks prior to recording.

### In vivo photometry data collection and analysis

Before each recording session, the background signal was measured by connecting the patch cable from the iFLiP setup to an unimplanted fiber cannula. The cannula tip was directed toward a black surface in a dark room to minimize reflected excitation light. Because the background signal depends on excitation intensity, a separate background recording was obtained for each laser power used in the experiment. Laser power was approximately 3–10 µW for head-fixed recordings and less than 1 µW for freely moving recordings.

For the head-fixed photometry experiments shown in Figure 5, mice were placed on a custom-built one-dimensional rotary treadmill with a 20-cm diameter wheel. The iFLiP recordings were performed with a sampling rate of 10 Hz. For enforced locomotion experiments, a motorized treadmill was driven at approximately 5 cm/s. Each session consisted of at least 10 min of baseline recording, 5 min of enforced running, and 15–20 min of post-running rest. Locomotor activity was measured using a rotary encoder (US Digital, MA3-A10-250-N) coupled to the treadmill axle. The encoder generated an analog voltage signal of approximately 0–5 V, which was digitized at 1 kHz using a National Instruments DAQ board (PCIe-6259), the MATLAB DAQ Toolbox, and custom MATLAB code. For the in vivo pharmacology experiments shown in Figures 4 and 5, mice first underwent one enforced running session. Drugs were then administered, followed by a 20– 30 min observation period and a second enforced running session after drug treatment. SKF83566 hydrobromide (Tocris) and eticlopride hydrochloride (Tocris) were dissolved in saline at a final concentration of 100 and 50 μg/mL, respectively.

For the freely moving open-field experiments shown in Figure 6, a two-channel iFLiP system, in which signals from two independent channels were detected simultaneously using a two-channel TCSPC board (TH260P-DUAL), was used to simultaneously record from two mice. For each recording, a mouse was connected to the optical patch cable through a pigtailed fiber-optic rotary joint (Doric) and placed in an open-top arena measuring 38.5 × 38.5 × 38.5 cm. The iFLiP recordings were performed with a sampling rate of 1 Hz. Mice were maintained on a standard 12-h light/dark cycle with ad libitum access to food and water. An overhead-mounted night-vision camera (ELP, ELP-USBFHD05MT-KL156IR) was used to record mouse position, and mouse motion was analyzed using ezTrack (Pennington et al., 2019). For freely moving pharmacology experiments, drugs were administered intraperitoneally 3 h after the light-to-dark or dark-to-light transition. Istradefylline (Tocris) and ACT-462206 (MedChemExpress) were dissolved in DMSO and then diluted in saline to a final concentration of 2.5 and 50 mg/mL, respectively. Scopolamine (Tocris) was dissolved in saline at a final concentration of 250 μg/mL.

### Data quantification and statistical analysis

All iFLiP data quantification was performed using custom MATLAB software. Because a reduction in MPET indicates an increase in PKA activities, the y-axes of MPET plots were flipped. Statistical tests were performed with MATLAB 2013b–2022a or GraphPad Prism 11. Data distribution was assessed using the Shapiro–Wilk test, and equality of variance was assessed using the F test or Brown–Forsythe test, as appropriate. Data that were normally and equally distributed were compared using a two-tailed paired or unpaired Student’s t-test or a one-way or two-way ANOVA with Bonferroni’s or Dunnett’s post hoc test. Data that were not normally or equally distributed were compared using the Mann-Whitney U test, Wilcoxon signed-rank test, Welch’s test, the Kruskal–Wallis test with Dunn’s post hoc test, the Friedman test with Dunn’s post hoc test, or Welch’s ANOVA test with Dunnett’s post hoc test. For the kinetics analysis of striatal tAKARα, the outliers were removed based on the ROUT method. All statistical results are in Table S2.

## Supplementary Figures and Legends

**Figure S1.**
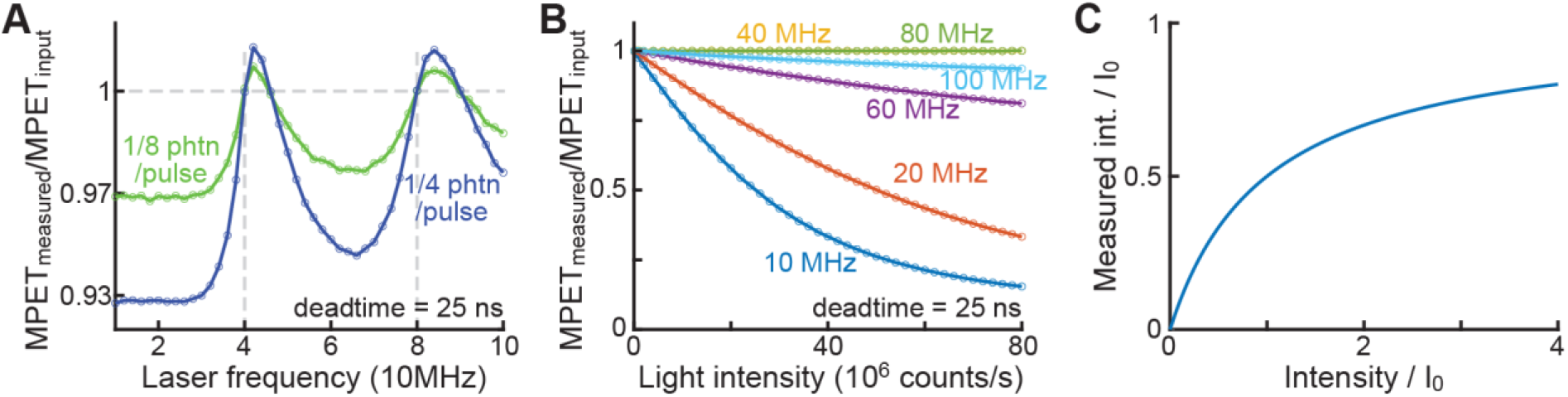
Simulation for the effect of TCSPC deadtime on intensity and lifetime measurements. Related to Figure 1. (A) For a given deadtime, error in lifetime (calculated using MPET) may occur except when the inter-pulse duration is a factor of deadtime (i.e., the product of laser frequency and deadtime gives a whole number). (B) For a 25 ns deadtime, most laser frequencies, but not 40 and 80 MHz, suffer intensity-dependent lifetime measurement errors. (C) The instrument deadtime limits the apparent photon counts (i.e., measured intensity). I_0_ = 1/deadtime. A practical system limit for incoming signals is arbitrarily set to I_0_, when 50% of the signal can be collected. For a 25 ns deadtime, this corresponds to the collection of 2×10^7^ photon/s. Phtn: photon

**Figure S2.**
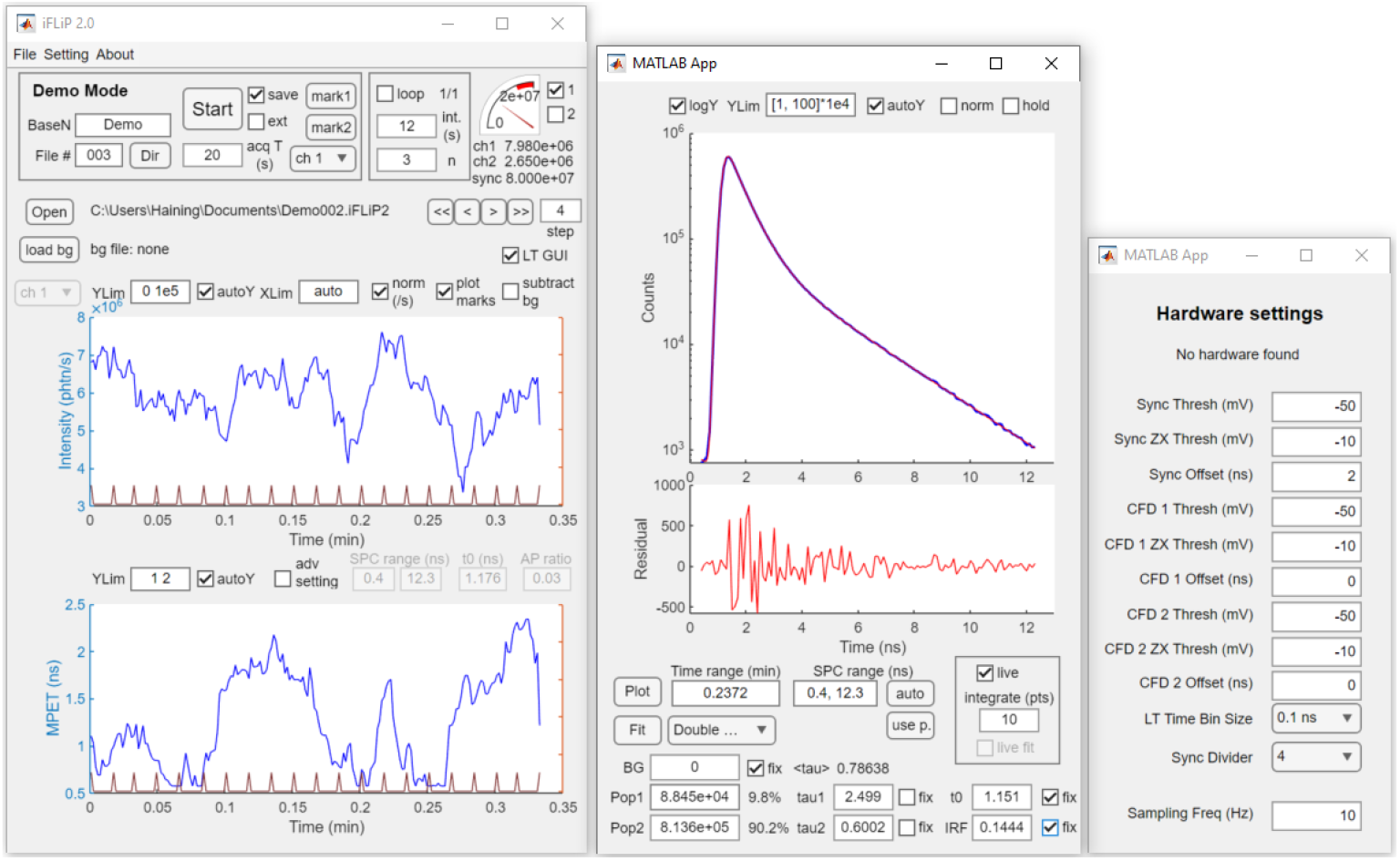
Software graphical user interface. Related to Figure 1.

**Figure S3.**
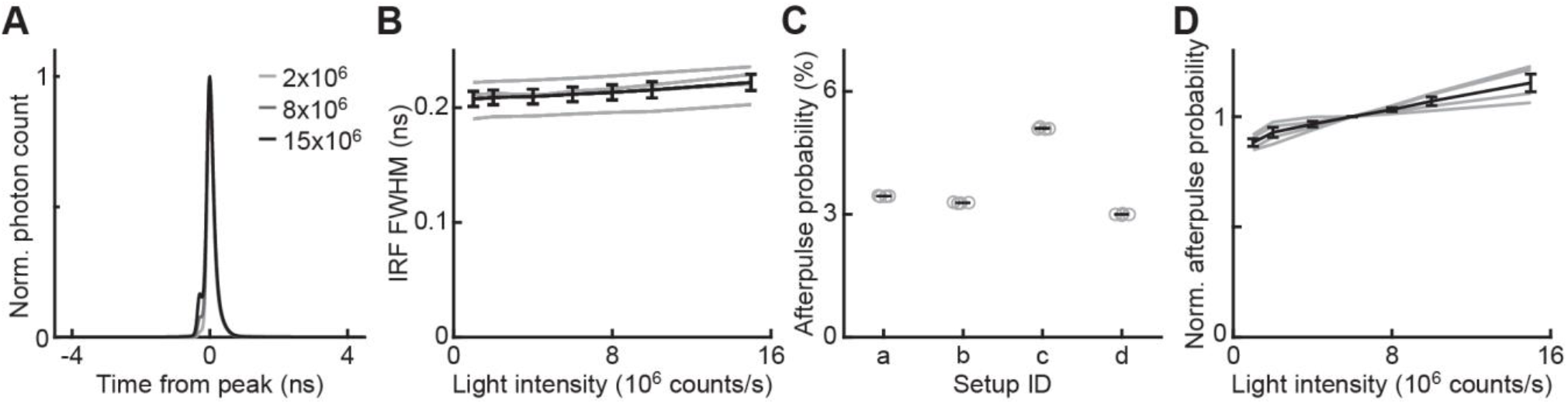
Example impulse response function of the iFLiP system. Related to Figure 1. (A) Example impulse response functions at 2×10^6^, 8×10^6^, and 15×10^6^ photon counts per second. (B) Mean (black) and individual (gray) full-width half-maximum of the impulse response function (n = 4 setups). (C) Mean (black line) and individual (gray dots) median afterpulse probability for all setups used (n = 4 measurements per setup). (D) Mean (black) and individual (gray) afterpulse intensity normalized to the median afterpulse intensity (n = 4 setups). All error bars: standard error.

**Figure S4.**
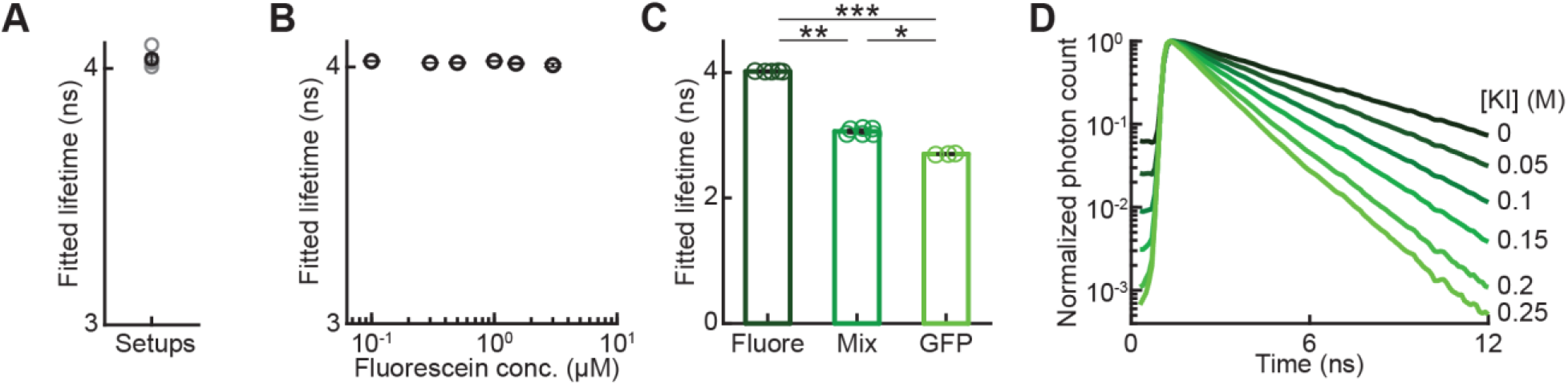
Additional data for in vitro validation of iFLiP. Related to Figure 2. (A) Fitted lifetime of fluorescein (1µM) across four setups. (B) Fitted lifetime of fluorescein across the indicated concentrations, corresponding to intensity groups shown in Fig. 2D. From left to right, n = 4, 4, 5, 8, 5, 5, respectively. (C) Fitted lifetime of fluorescein (0.5 µM, n = 5), GFP (0.5 µM, n = 3), and a mixture (n = 6) of fluorescein (0.125 µM) and GFP (0.375 µM), from the same samples plotted in Figure 2D. (D) Normalized lifetime curve of fluorescein (1 µM) quenched by different concentrations of KI. n = 8 for [KI] = 0 M; n = 3 for all other concentrations. All error bars: standard error. *: p < 0.05; **: p < 0.01; ***: p < 0.001.

**Figure S5.**
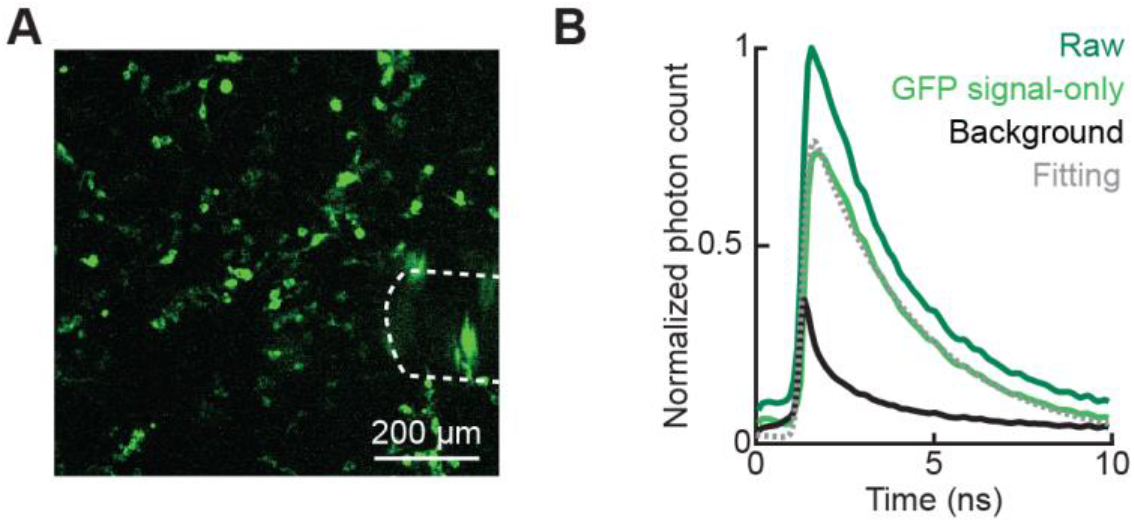
In vitro validation of lifetime measurement in HEK cells. Related to Figure 3. (A) Example image of tAKARβ expression (green) in HEK cells. Dotted line: location of the fiber. (B) Example normalized photon counts of raw GFP expression in HEK cells, GFP after background subtraction (signal-only), and background. Dotted line: fitting of the background-subtracted GFP signal.

**Figure S6.**
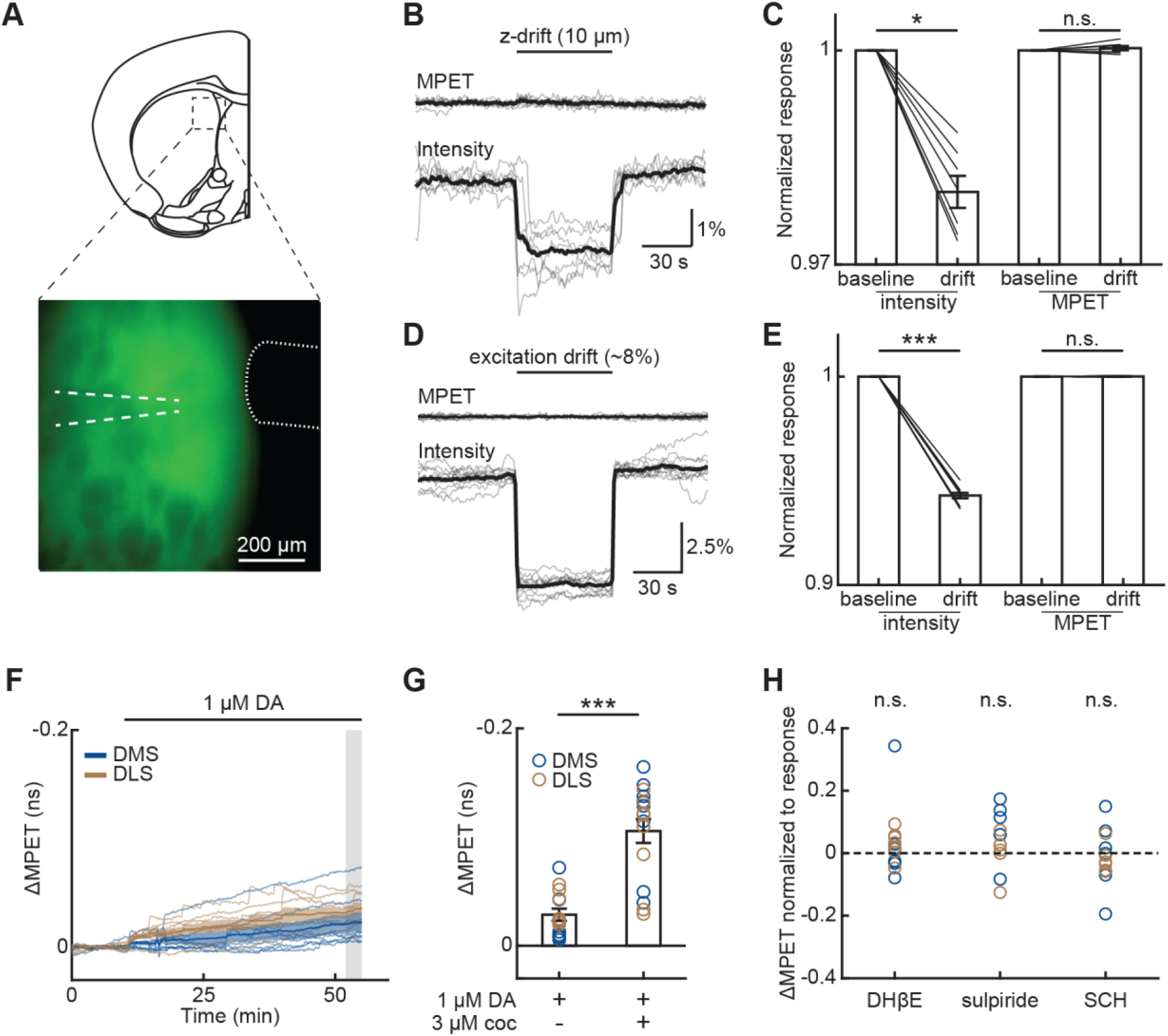
Ex vivo validation of lifetime measurement stability and sensitivity to dopamine. Related to Figure 3. (A) Example image of cd-tAKARα expression (green) in a striatal slice. Dotted line: location of the fiber. Dashed line: the stimulation electrode. (B–E) Intensity change induced by fiber z-depth adjustment (B and C, n = 7; also see STAR Methods) or excitation adjustment (D and E, n = 11). (F) ΔMPET of cd-tAKARα expressed in DMS (blue, n = 7) and DLS (gold, n = 7) slices in response to 1 µM dopamine (DA). Plotted in the same scale as Figure 3H. Shaded rectangles: analysis time window for panel (K). (G) Quantification of ΔMPET in response to dopamine (1 µM) alone (also see panel F) and to dopamine (DA, 1 µM) followed by cocaine (coc, 3 µM, n = 8 for both) (also see Figure 3H). +: with; –: without; blue dots: DMS datapoints; gold dots: DLS datapoints. (H) Baseline ΔMPET in response to DHβE (1µM), sulpiride (5 µM), or SCH (100 nM) in addition to sulpiride, normalized to evoked response before drug administration. DA: dopamine; coc: cocaine; +: with; –: without. Throughout the figure, mean (darker lines) and s.e.m. (shaded area and error bars) are shown. Thin lines are individual recordings. n.s.: not significant, *: p < 0.05, **: p < 0.01, ***: p < 0.001. See Table S2 for detailed statistics.

**Figure S7.**
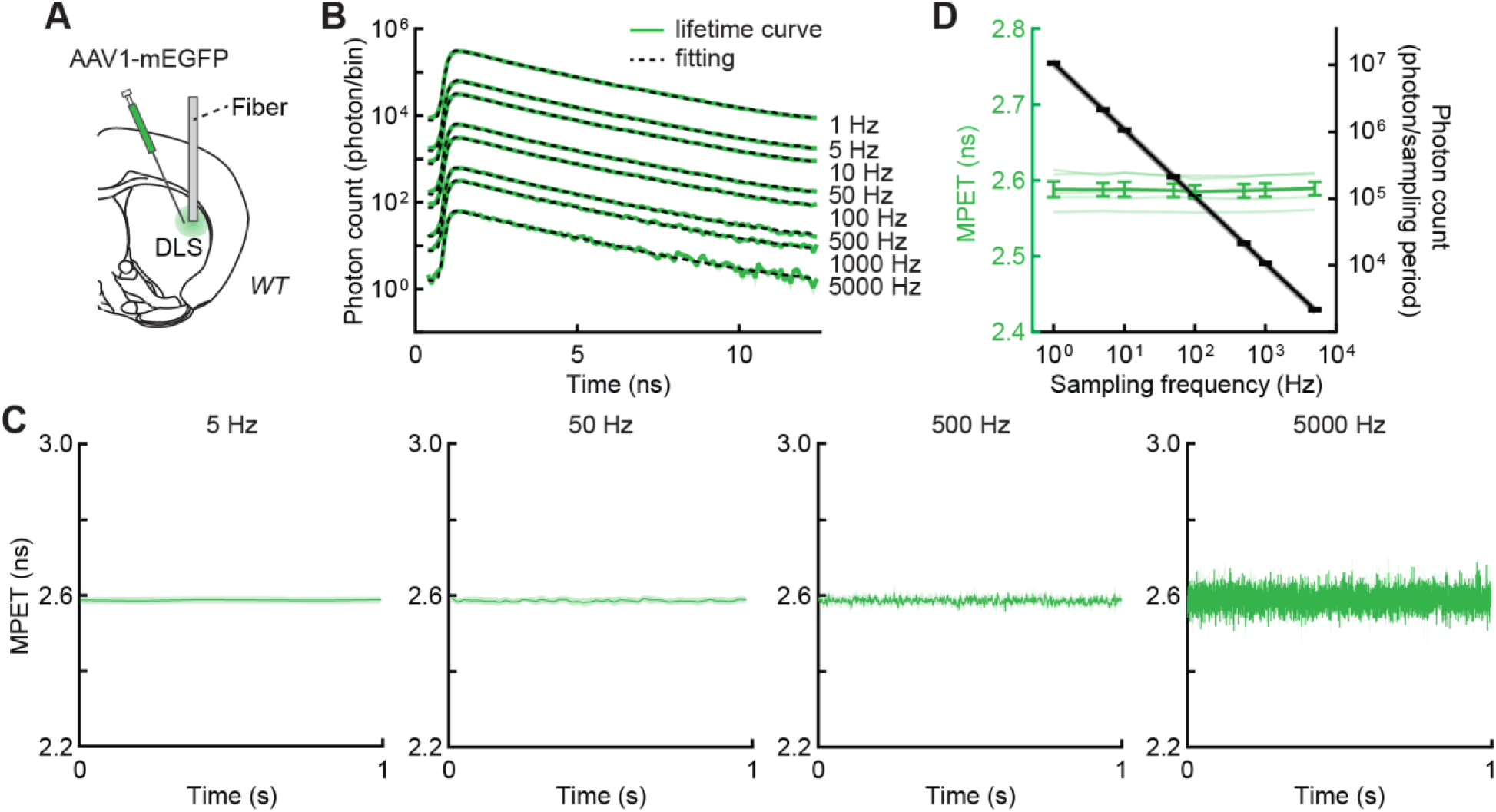
In vivo iFLiP can report GFP lifetime at different sampling frequencies. Related to Figure 4. (A) Schematics of virus injections and fiber cannula implantation. (B) Example average traces of 10 lifetime curves recording GFP with 1–5000 Hz sampling frequencies. Double exponential fitting curves are shown as black dashed lines. (C) Averaged traces of GFP MPET with multiple sampling frequencies (n = 5). (D) MPET and photon count across different sampling frequencies (n = 5). Throughout, mean (darker lines) and s.e.m. (shaded area and error bars) are shown.

**Figure S8.**
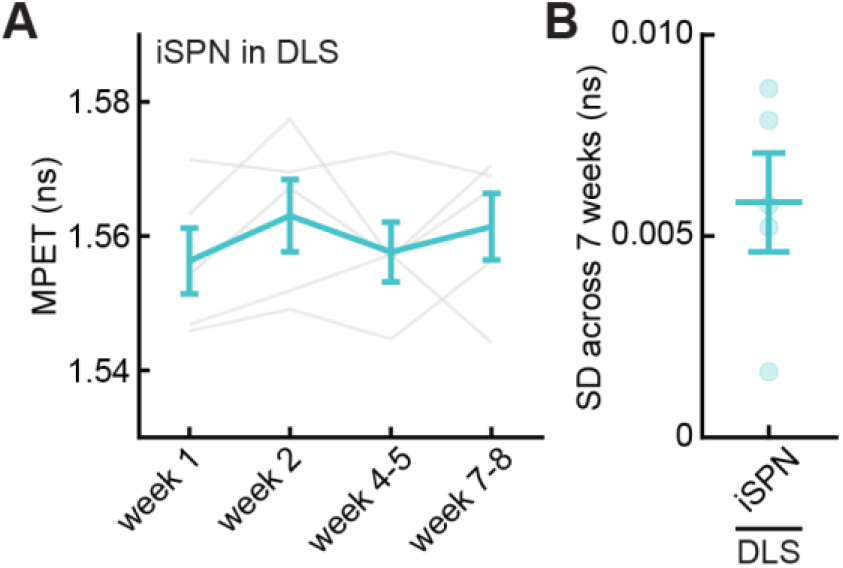
iFLiP enables stable measurements of baseline across weeks. Related to Figure 5. (A, B) Time course (A) and standard deviation (B) of repeated measurements of the same animal across time. n = 5.

**Figure S9.**
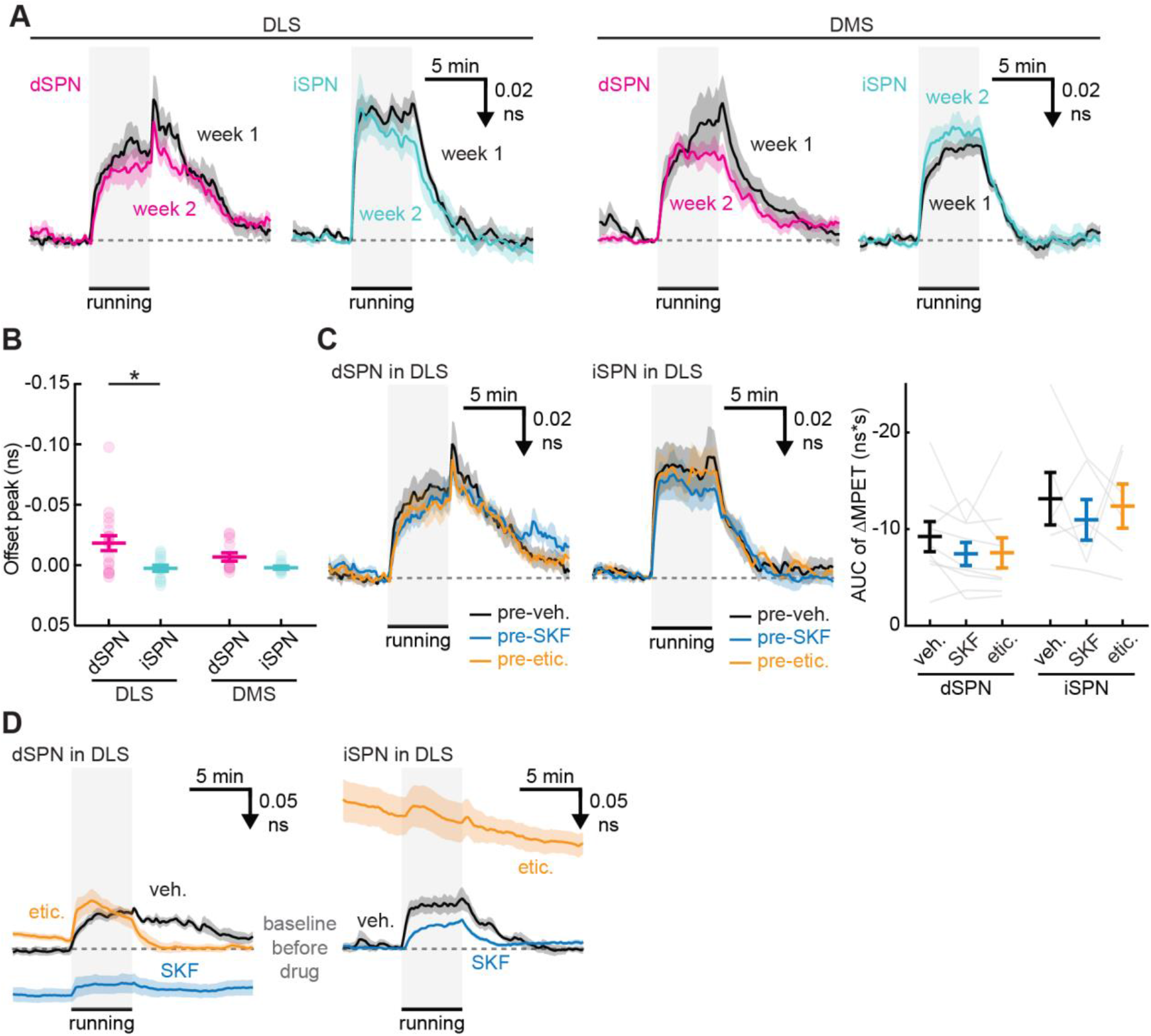
iFLiP detects the enforced running-induced PKA activity across weeks and the pharmacological effect on it. Related to Figure 5. (A) Averaged traces of enforced running-induced MPET changes of tAKARα across weeks in dSPNs (black: week 1; magenta: week 2) and iSPNs (black: week 1; cyan: week 2) of the DLS, and tAKARα in dSPNs (black: week 1; magenta: week 2) and iSPNs (black: week 1; cyan: week 2) of the DMS. (B) Peak amplitudes 5 min during enforced running, and 0.5–1 min after enforced running were quantified by normalizing to the offset of enforced running. Sample sizes were n (bout/mouse) = 18/9, 14/7, 12/6, and 12/6 for DLS dSPNs, DLS iSPNs, DMS dSPNs, and DMS iSPNs, respectively. (C) Averaged traces and quantification of enforced running-induced MPET changes of tAKARα in dSPNs and iSPNs of the DLS before drug injections. n = 9 and 7 for DLS dSPNs and DLS iSPNs, respectively. (D) Pharmacological effects on enforced running-induced PKA changes of dSPN in the DLS and iSPN. The MPET change is calculated from the basal MPET before drug injections. Veh.: vehicle; SKF: SKF83566; etic.: eticlopride. Throughout the figure, mean (darker lines) and s.e.m. (shaded area and error bars) are shown. *: p < 0.05, **: p < 0.01, ***: p < 0.001. See Table S2 for detailed statistics.

**Figure S10.**
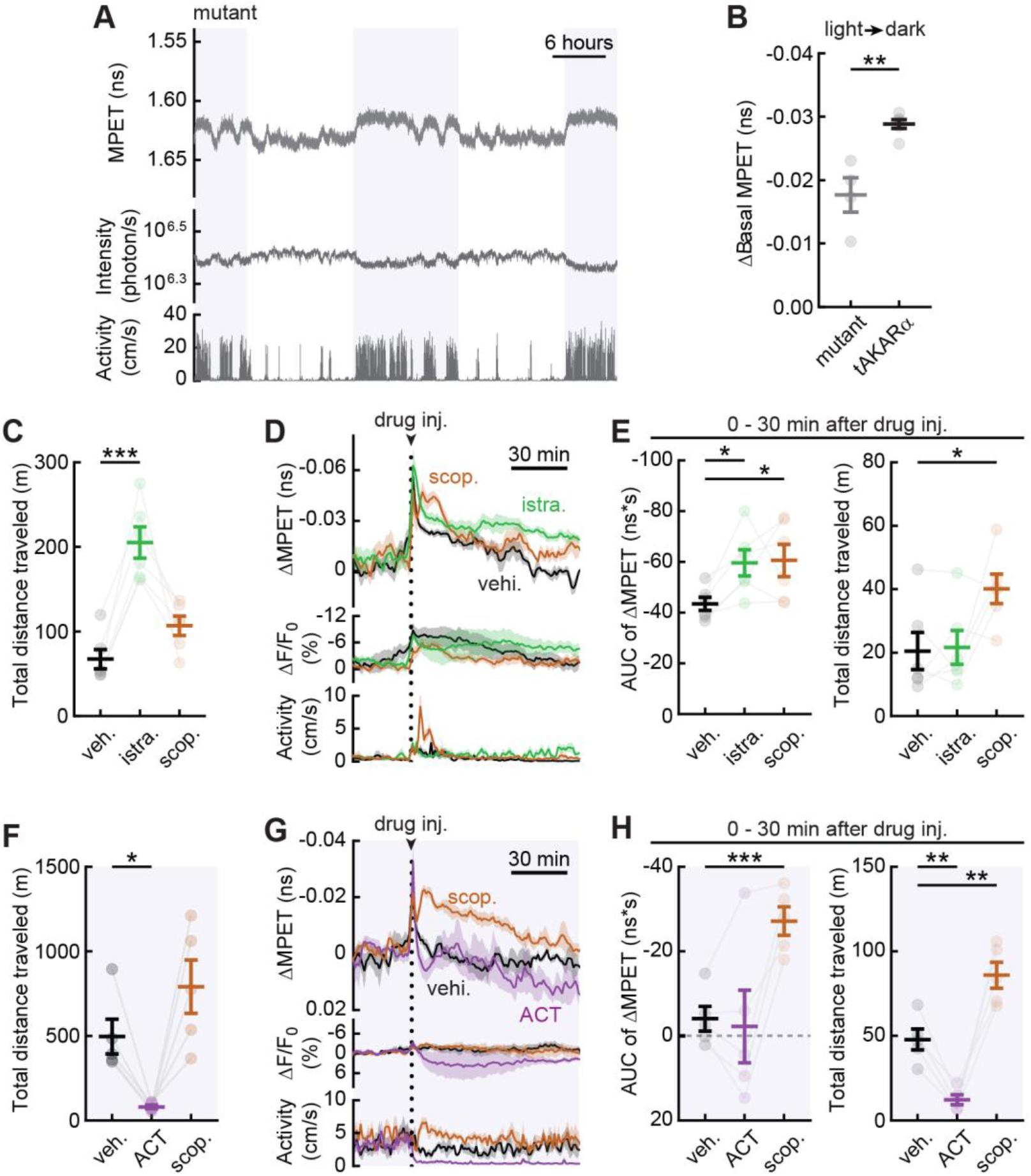
Additional analyses of basal forebrain PKA dynamics and pharmacological manipulations. Related to Figure 6. (A) Representative traces of a 48-hour recording of tAKARα in the basal forebrain. (B) Quantification of the baseline change between the light and dark phases with mutant and tAKARα. n = 4 and 6 for mutant and tAKARα, respectively. (C) Quantification of the total distance traveled during the 0 – 5 hours following drug injections. n = 6; related to Figures 6K and L. (D, E) Expanded view of Figure 6K during 0–1.5 hours following drug injections, and quantification of the AUC for ΔMPET and total distance traveled during 0–30 minutes following drug injections. n = 6. (F) Quantification of the total distance traveled during 0–5 hours following drug injections. n = 5. (G, H) Expanded view of Figure 6M during 0–1.5 hours following drug injections, and quantification of the AUC for ΔMPET and total distance traveled during 0–30 minutes following drug injections. n = 5. Veh: vehicle; istra.: istradefylline; scop.: scopolamine; ACT: ACT-462206. Throughout the figure, mean (darker lines) and s.e.m. (shaded area and error bars) are shown; darker and white backgrounds indicate the dark and light phases, respectively. *: p < 0.05, **: p < 0.01, ***: p < 0.001. See Table S2 for detailed statistics.

## REFERENCES

1. Sabatini, B.L., and Tian, L. (2020). Imaging Neurotransmitter and Neuromodulator Dynamics In Vivo with Genetically Encoded Indicators, 10.1016/j.neuron.2020.09.036.

2. Day-Cooney, J., Dalangin, R., Zhong, H., and Mao, T. (2022). Genetically encoded fluorescent sensors for imaging neuronal dynamics in vivo, 10.1111/jnc.15608.

3. Yang, Y., Li, B., and Li, Y. (2024). Genetically Encoded Sensors for the In Vivo Detection of Neurochemical Dynamics. Annu. Rev. Anal. Chem. 17. 10.1146/annurev-anchem-061522-044819.

4. Yasuda, R. (2006). Imaging spatiotemporal dynamics of neuronal signaling using fluorescence resonance energy transfer and fluorescence lifetime imaging microscopy, 10.1016/j.conb.2006.08.012.

5. Yellen, G., and Mongeon, R. (2015). Quantitative two-photon imaging of fluorescent biosensors at Elsevier Ltd, 10.1016/j.cbpa.2015.05.024.

6. Berezin, M.Y., and Achilefu, S. (2010). Fluorescence lifetime measurements and biological imaging. Chem. Rev. 110. 10.1021/cr900343z.

7. Yasuda, R., Harvey, C.D., Zhong, H., Sobczyk, A., Van Aelst, L., and Svoboda, K. (2006). Supersensitive Ras activation in dendrites and spines revealed by two-photon fluorescence lifetime imaging. Nat. Neurosci. 9, 283–291. 10.1038/nn1635.

8. Díaz-García, C.M., Mongeon, R., Lahmann, C., Koveal, D., Zucker, H., and Yellen, G. (2017). Neuronal Stimulation Triggers Neuronal Glycolysis and Not Lactate Uptake. Cell Metab. 26. 10.1016/j.cmet.2017.06.021.

9. Ma, L., Jongbloets, B.C., Xiong, W.H., Melander, J.B., Qin, M., Lameyer, T.J., Harrison, M.F., Zemelman, B. V., Mao, T., and Zhong, H. (2018). A Highly Sensitive A-Kinase Activity Reporter for Imaging Neuromodulatory Events in Awake Mice. Neuron 99, 665–679.e5. 10.1016/j.neuron.2018.07.020.

10. Lee, S.J., Lodder, B., Chen, Y., Patriarchi, T., Tian, L., and Sabatini, B.L. (2021). Cell-type-specific asynchronous modulation of PKA by dopamine in learning. Nature 590, 451–456. 10.1038/s41586-020-03050-5.

11. Ma, L., Day-Cooney, J., Benavides, O.J., Muniak, M.A., Qin, M., Ding, J.B., Mao, T., and Zhong, H. (2022). Locomotion activates PKA through dopamine and adenosine in striatal neurons. Nature 611, 762–768. 10.1038/s41586-022-05407-4.

12. Yahiro, T., Bayless-Edwards, L., Jones, J.A., Ma, L., Qin, M., Mao, T., and Zhong, H. (2025). A high-performance genetically encoded sensor for cellular imaging of PKC activity in vivo. Nat. Commun. 16:6378, 1–12.

13. Massengill, C.I., Bayless-Edwards, L., Ceballos, C.C., Cebul, E.R., Cahill, J., Bharadwaj, A., Wilson, E., Qin, M., Whorton, M.R., Baconguis, I., et al. (2022). Sensitive genetically encoded sensors for population and subcellular imaging of cAMP in vivo. Nat. Methods 19, 1461–1471. 10.1038/s41592-022-01646-5.

14. Zhang, S.X., Kim, A., Madara, J.C., Zhu, P.K., Christenson, L.F., Lutas, A., Kalugin, P.N., Sunkavalli, P.S., Jin, Y., Pal, A., et al. (2025). Stochastic neuropeptide signals compete to calibrate the rate of satiation. Nature 637. 10.1038/s41586-024-08164-8.

15. Lodder, B., Kamath, T., Savenco, E., Röring, B., Siegel, M., Chouinard, J.A., Lee, S.J., Zagoren, C., Rosen, P., Hartman, I., et al. (2025). Absolute measurement of fast and slow neuronal signals with fluorescence lifetime photometry at high temporal resolution. Neuron 113. 10.1016/j.neuron.2025.08.013.

16. Koveal, D., Rosen, P.C., Meyer, D.J., Díaz-García, C.M., Wang, Y., Cai, L.H., Chou, P.J., Weitz, D.A., and Yellen, G. (2022). A high-throughput multiparameter screen for accelerated development and optimization of soluble genetically encoded fluorescent biosensors. Nat. Commun. 13. 10.1038/s41467-022-30685-x.

17. van der Linden, F.H., Mahlandt, E.K., Arts, J.J.G., Beumer, J., Puschhof, J., de Man, S.M.A., Chertkova, A.O., Ponsioen, B., Clevers, H., van Buul, J.D., et al. (2021). A turquoise fluorescence lifetime-based biosensor for quantitative imaging of intracellular calcium. Nat. Commun. 12. 10.1038/s41467-021-27249-w.

18. Laviv, T., Scholl, B., Parra-Bueno, P., Foote, B., Zhang, C., Yan, L., Hayano, Y., Chu, J., and Yasuda, R. (2020). In Vivo Imaging of the Coupling between Neuronal and CREB Activity in the Mouse Brain. Neuron 105. 10.1016/j.neuron.2019.11.028.

19. Lee, S.J., Chen, Y., Lodder, B., and Sabatini, B.L. (2019). Monitoring behaviorally induced biochemical changes using fluorescence lifetime photometry. Front. Genet. 10. 10.3389/fnins.2019.00766.

20. Gadella, T.W.J., Jovin, T.M., and Clegg, R.M. (1993). Fluorescence lifetime imaging microscopy (FLIM): Spatial resolution of microstructures on the nanosecond time scale. Biophys. Chem. 48. 10.1016/0301-4622(93)85012-7.

21. Digman, M.A., Caiolfa, V.R., Zamai, M., and Gratton, E. (2008). The phasor approach to fluorescence lifetime imaging analysis. Biophys. J. 94. 10.1529/biophysj.107.120154.

22. Barber, P.R., Ameer-Beg, S.M., Gilbey, J., Carlin, L.M., Keppler, M., Ng, T.C., and Vojnovic, B. (2009). Multiphoton time-domain fluorescence lifetime imaging microscopy: Practical application to protein-protein interactions using global analysis. In Journal of the Royal Society Interface 10.1098/rsif.2008.0451.focus.

23. Torbeck, R.L., Schilling, L., Khorasani, H., Dover, J.S., Arndt, K.A., and Saedi, N. (2019). Evolution of the Picosecond Laser: A Review of Literature, 10.1097/DSS.0000000000001697.

24. Gedcke, D.A., and McDonald, W.J. (1967). A constant fraction of pulse height trigger for optimum time resolution, 10.1016/0029-554X(67)90145-0 https://doi.org/10.1016/0029-554X(67)90145-0.

25. Coates, P.B. (1968). The correction for photon “pile-up” in the measurement of radiative lifetimes. J. Phys. E. 1. 10.1088/0022-3735/1/8/437.

26. Salthammer, T. (1992). Numerical simulation of pile-up distorted time-correlated single photon counting (TCSPC) data. J. Fluoresc. 2. 10.1007/BF00866385.

27. Klonis, N., and Sawyer, W.H. (1996). Spectral properties of the prototropic forms of fluorescein in aqueous solution. J. Fluoresc. 6. 10.1007/BF00732054.

28. Zhang, X.F., Zhang, J., and Liu, L. (2014). Fluorescence properties of twenty fluorescein derivatives: Lifetime, quantum yield, absorption and emission spectra. J. Fluoresc. 24. 10.1007/s10895-014-1356-5.

29. Hammer, M., Schweitzer, D., Richter, S., and Königsdörffer, E. (2005). Sodium fluorescein as a retinal pH indicator? Physiol. Meas. 26. 10.1088/0967-3334/26/4/N01.

30. Watt, R.M., and Voss, E.W. (1979). Solvent perturbation of the fluorescence of fluorescein bound to specific antibody. Fluorescence quenching of the bound fluorophore by iodide. J. Biol. Chem. 254. 10.1016/s0021-9258(17)37827-4.

31. Koenig, M.H., Yi, E.P., Sandridge, M.J., Mathew, A.S., and Demas, J.N. (2015). Open-box approach to measuring fluorescence quenching using an ipad screen and digital SLR camera. J. Chem. Educ. 92. 10.1021/ed500373d.

32. Chen, Y., Saulnier, J.L., Yellen, G., and Sabatini, B.L. (2014). A PKA activity sensor for quantitative analysis of endogenous GPCR signaling via 2-photon FRET-FLIM imaging. Front. Pharmacol. 5 *APR*, 56. 10.3389/fphar.2014.00056.

33. Surmeier, D.J., Plotkin, J., and Shen, W. (2009). Dopamine and synaptic plasticity in dorsal striatal circuits controlling action selection, 10.1016/j.conb.2009.10.003.

34. Tritsch, N.X., and Sabatini, B.L. (2012). Dopaminergic Modulation of Synaptic Transmission in Cortex and Striatum, 10.1016/j.neuron.2012.09.023.

35. Klaus, A., Alves Da Silva, J., and Costa, R.M. (2019). What, If, and When to Move: Basal Ganglia Circuits and Self-Paced Action Initiation, 10.1146/annurev-neuro-072116-031033.

36. Lovinger, D.M. (2010). Neurotransmitter roles in synaptic modulation, plasticity and learning in the dorsal striatum, 10.1016/j.neuropharm.2010.01.008.

37. Gerfen, C.R., and Surmeier, D.J. (2011). Modulation of striatal projection systems by dopamine. Annu. Rev. Neurosci. 34, 441–466. 10.1146/annurev-neuro-061010-113641.

38. Threlfell, S., Lalic, T., Platt, N.J., Jennings, K.A., Deisseroth, K., and Cragg, S.J. (2012). Striatal dopamine release is triggered by synchronized activity in cholinergic interneurons. Neuron 75, 58–64. 10.1016/j.neuron.2012.04.038.

39. Kramer, P.F., Brill-Weil, S.G., Cummins, A.C., Zhang, R., Camacho-Hernandez, G.A., Newman, A.H., Eldridge, M.A.G., Averbeck, B.B., and Khaliq, Z.M. (2022). Synaptic-like axo-axonal transmission from striatal cholinergic interneurons onto dopaminergic fibers. Neuron 110. 10.1016/j.neuron.2022.07.011.

40. Liu, C., Cai, X., Ritzau-Jost, A., Kramer, P.F., Li, Y., Khaliq, Z.M., Hallermann, S., and Kaeser, P.S. (2022). An action potential initiation mechanism in distal axons for the control of dopamine release. Science (80-.). 375. 10.1126/science.abn0532.

41. Gerfen, C.R., Engber, T.M., Mahan, L.C., Susel, Z., Chase, T.N., Monsma, F.J., and Sibley, D.R. (1990). D1 and D2 dopamine receptor-regulated gene expression of striatonigral and striatopallidal neurons. Science (80-.). 250, 1429–1432. 10.1126/science.2147780.

42. Li, W., Ma, L., Yang, G., and Gan, W.B. (2017). REM sleep selectively prunes and maintains new synapses in development and learning. Nat. Neurosci. 20, 427–437. 10.1038/nn.4479.

43. Paukert, M., Agarwal, A., Cha, J., Doze, V.A., Kang, J.U., and Bergles, D.E. (2014). Norepinephrine controls astroglial responsiveness to local circuit activity. Neuron 82, 1263–1270. 10.1016/j.neuron.2014.04.038.

44. Lakowicz, J.R. (2006). Principles of fluorescence spectroscopy 10.1007/978-0-387-46312-4.

45. Silva, P.L., Savchuk, O.A., Gallo, J., García-Hevia, L., Bañobre-López, M., and Nieder, J.B. (2020). Mapping intracellular thermal response of cancer cells to magnetic hyperthermia treatment. Nanoscale 12. 10.1039/c9nr10370h.

46. Kamath, T., Lodder, B., Bilsel, E., Green, I., Dalangin, R., Raghubardayal, M., Wang, W., Capelli, P., Legister, J., Timmins, J., et al. (2025). Hunger modulates exploration through suppression of dopamine signaling in the tail of the striatum. Neuron 113. 10.1016/j.neuron.2025.09.009.

47. Fink, R., Imai, S., Gockel, N., Lauer, G., Renken, K., Wietek, J., Lamothe-Molina, P.J., Fuhrmann, F., Mittag, M., Ziebarth, T., et al. (2025). PinkyCaMP a mScarlet-based calcium sensor with exceptional brightness, photostability, and multiplexing capabilities. bioRxiv.

48. Mohebi, A., Wei, W., Pelattini, L., Kim, K., and Berke, J.D. (2024). Dopamine transients follow a striatal gradient of reward time horizons. Nat. Neurosci. 27. 10.1038/s41593-023-01566-3.

49. Adrover, M.F., Shin, J.H., Quiroz, C., Ferré, S., Lemos, J.C., and Alvarez, V.A. (2020). Prefrontal cortex-driven dopamine signals in the striatum show unique spatial and pharmacological properties. J. Neurosci. 40. 10.1523/JNEUROSCI.1327-20.2020.

50. Dong, Y., Li, Y., Xiang, X., Xiao, Z.C., Hu, J., Li, Y., Li, H., and Hu, H. (2023). Stress relief as a natural resilience mechanism against depression-like behaviors. Neuron 111. 10.1016/j.neuron.2023.09.004.

51. Hunnicutt, B.J., Long, B.R., Kusefoglu, D., Gertz, K.J., Zhong, H., and Mao, T. (2014). A comprehensive thalamocortical projection map at the mesoscopic level. Nat. Neurosci. 17, 1276–1285. 10.1038/nn.3780.

52. Meng, C., Zhou, J., Papaneri, A., Peddada, T., Xu, K., and Cui, G. (2018). Spectrally Resolved Fiber Photometry for Multi-component Analysis of Brain Circuits. Neuron 98, 707–717.e4. 10.1016/j.neuron.2018.04.012.

